# Escaping the drug-bias trap: using debiasing design to improve interpretability and generalization of drug-target interaction prediction

**DOI:** 10.1101/2024.09.12.612771

**Authors:** Pei-Dong Zhang, Jianzhu Ma, Ting Chen

## Abstract

Considering the high cost associated with determining reaction affinities through in-vitro experiments, virtual screening of potential drugs bound with specific protein pockets from vast compounds is critical in AI-assisted drug discovery. Deep-leaning approaches have been proposed for Drug-Target Interaction (DTI) prediction. However, they have shown overestimated accuracy because of the drug-bias trap, a challenge that results from excessive reliance on the drug branch in the traditional drug-protein dual-branch network approach. This casts doubt on the interpretability and generalizability of existing Drug-Target Interaction (DTI) models. Therefore, we introduce UdanDTI, an innovative deep-learning architecture designed specifically for predicting drug-protein interactions. UdanDTI applies an unbalanced dual-branch system and an attentive aggregation module to enhance interpretability from a biological perspective. Across various public datasets, UdanDTI demonstrates outstanding performance, outperforming state-of-the-art models under in-domain, cross-domain, and structural interpretability settings. Notably, it demonstrates exceptional accuracy in predicting drug responses of two crucial subgroups of Epidermal Growth Factor Receptor (EGFR) mutations associated with non-small cell lung cancer, consistent with experimental results. Meanwhile, UdanDTI could complement the advanced molecular docking software DiffDock. The codes and datasets of UdanDTI are available at https://github.com/CQ-zhang-2016/UdanDTI.

## 1 INTRODUCTION

Identifying drug-target interactions represents a pivotal quest in drug discovery to uncover bioactive compound candidates and their potential interactions with target proteins. While traditional in-vitro experimentation remains the gold standard for evaluating drug-target binding affinities [1], its utility is often limited by its exorbitant costs, time-consuming nature, and inefficiencies [2]. Consequently, computer-based virtual screening has emerged as a promising alternative for rapid Drug-Target Interaction (DTI) prediction and elucidation of binding patterns.

Molecular docking is a physics-based virtual screening technique, but the expansion of molecular space renders docking every potential drug-target pair impractical. For instance, using commercial software to dock 10 billion candidate drug molecules would require approximately 30 years [3]. Conversely, machine learning enables DTI models to quickly predict potential affinity from structure-independent raw features [4-11]. The most popular dual-branch network model encodes proteins and drugs separately using two branches and then merges them for interaction predictions. In the dual-branch deep learning-based architectures, researchers have explored various encoding modules like Convolutional Neural Networks (CNNs) [12-16], Graph Neural Networks (GNNs) [17-22], Deep Neural Networks (DNNs) [23, 24], and Transformers [25-27], utilizing 1D protein sequences and 2D drug representations to decipher binding patterns. Additionally, incorporating multi-modal information has also improved performance [28, 29]. Recently, Large Language Models (LLMs) in bioinformatics have been introduced to support DTI prediction by encoding molecular properties [30-32].

Despite these advancement of deep learning methods, generalizability and interpretability remain two significant challenges [15, 33]. Potential series bias in data distribution is the main reason. Earlier work has highlighted this bias using methods such as modeling interactions [25], focusing ligand [26], and introducing void protein experiments [22]. In fact, ligand bias is amplified in dual-branch network models with inadequately designed interactive modules [22, 26, 34]. Eid et al. demonstrated that existing interaction predictors might separately predict the binding degree ratio of samples on each branch from training data and then multiply these predictions together to obtain the final prediction [35]. In the DTI field, this bias often stems from the simplicity of drug structures, making models prone to identifying drug-centric interaction sites while neglecting the complex spatial properties of protein pockets [22, 26]. We call this a “drug-bias trap”. The prevalent issue of the drug-bias trap is observed in models that overly rely on information from the training set or fall into overfitting due to high protein homology [15, 16, 25, 36]. Further investigations into drug-bias trap have been detailed in section 4 of the Supplementary Information, following the methodology outlined in [22].

To address these enduring challenges, we propose UdanDTI, an unbalanced dual-branch neural network, as a solution to enhance interpretability and generalization in DTI predictions (as illustrated in Fig.1a) of virtual screening. Distinct from previous balanced networks, UdanDTI considers disparate biochemical differences for proteins and drugs to prioritize unbalanced depths to ensure equitable attention to both branches. Specifically, UdanDTI uses protein sequences and Simplified Molecular Input Line Entry System (SMILES) strings as input. UdanDTI first captures a high-dimensional representation of the protein through LLMs like ESM [37] or ProtBert [38] with frozen weights, which encodes a wider range of contextual knowledge from the amino acid sequence. Meanwhile, the drug molecule is characterized as a graph, with atoms as nodes and atomic bonds as edges, and then a GCN-based module is used to encode local features of the drug spatial structure. Our novel aggregation module, depicted in Fig.1b, is specifically designed to capture mutual information effectively. Firstly, two parallel transformer layers are used to assimilate self-information of the drug and the protein, respectively. Two cross-attention layers consider the different modeling scales and facilitate the detection of local interaction patterns through cross-attention weights. The aggregation module ensures that features in every channel of the final output come from both branches, explicitly disrupting the independence of the branches and forcing the model to learn from the fused features. In the end, UdanDTI is augmented with a model called Maximum Classifier Discrepancy [39] (MCD, as shown in Fig.1c) to facilitate cross-domain adaptation, enhancing robustness and partly alleviating cross-domain interpretability issues.

**Fig 1.**
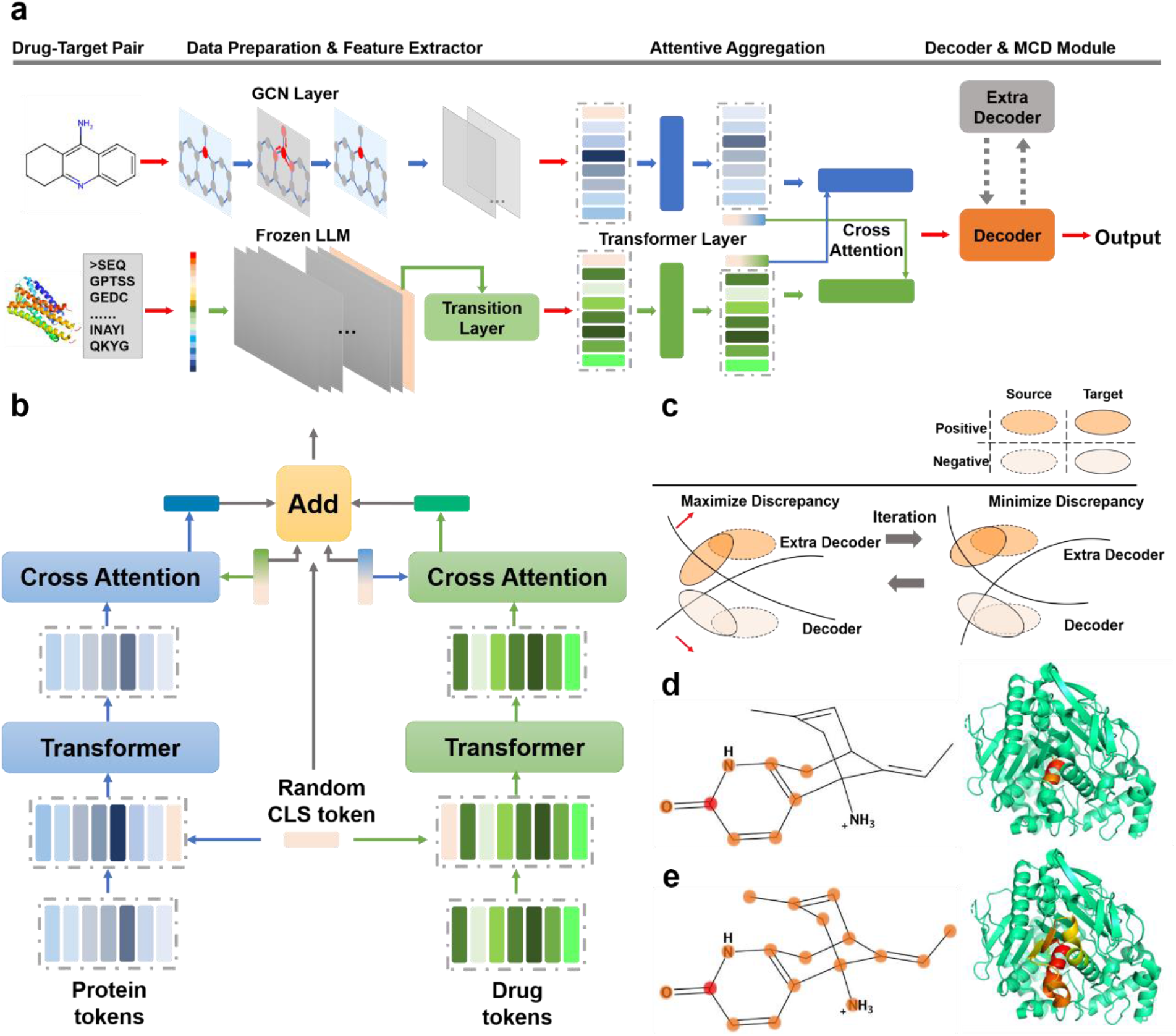
(a) The overall architecture of UdanDTI. A sampled drug-target pair, represented as a drug 3-D structure and a protein 1-D sequence, sequentially passes through four modules: Data Preparation, Feature Extractor, Attentive Aggregation, and Decoder. If the feature distributions of the test and training data differ significantly, an MCD module is activated with an additional decoder for unsupervised domain adaptation. The final output is a natural number ranging from 0 to 1, indicating the binding potential between the pair. (b) Details of the Attentive Aggregation module. It involves two parallel branches containing a concatenated transformer-cross attention layer. A randomly generated token, incorporated as a head, is added to the representations of the protein and the drug for transformer training. Then, the head tokens from both branches are exchanged to guide the crossattention training. Five tokens are weighted and summed to smooth the prediction. (c) Concept of the MCD module[43]. Two decoders (hyperplanes for decision-making) are trained to increase discrepancy in target domain data. Subsequently, the front-end network is trained to diminish discrepancy. This adversarial approach pulls target domain samples near the decision boundary into the correct class space. (d) and (e) are illustrations of an unbalanced dual-branch for protein Acetylcholinesterase and drug Huperzine A. (d) A 3-layer network’s receptive field adequately captures the crucial functional group pyridine-carbonyl (interaction functional group in Complex 1GPK). However, it only covers a small portion of the acetylcholinesterase helix. (e) A 24-layer network (the averaged depth of LLMs) effectively focuses on the spatial characteristics of the protein pocket but obscures the crucial nodes of the drug molecule.

Comparative evaluations against current state-of-the-art (SOTA) methods across in-domain, cross-domain, and interpretability metrics on independent public datasets confirm UdanDTI’s superiority. Moreover, UdanDTI performs well in two main subgroups (on the basis of sensitivity and structural changes) of epidermal growth factor receptor (EGFR) mutations. Besides, experiments indicate that UdanDTI could also be used as a reliable auxiliary tool in molecular docking tasks.

## 2 EVALUATION STRATEGIES AND METRICS

This study applies the proposed model across three public datasets, BindingDB [40], BioSNAP [41], and Human [42], to conduct a comparative analysis against seven state-of-the-art models. Additionally, drug responses to EGFR mutations [43] are also used as a test set to evaluate the performance of UdanDTI. Detailed description of baseline models and datasets can be found in Supplementary Information. We conduct two types of experiments: in-domain and cross-domain. Random splitting and cold-pair splitting strategies are used in in-domain experiments. The random splitting strategy divides the dataset into the training, validation, and testing sets at a ratio of 7:2:1. The cold-pair method reserves 5% and 10% DTI pairs to the validation and test sets, respectively, and removes all relevant drugs and proteins from the training set.

In cross-domain experiments, we apply a stricter clus-ter-based splitting strategy to split latent feature clusters of proteins and drugs in the testing set from those in the training set [15]. Furthermore, we partitioned BioSNAP and BindingDB using three other cluster-based splitting methods: with unseen drugs, where the drugs in the testing set are not present in the training set; one with unseen proteins, where the proteins in the testing set are not present in the training set; and one using sphere exclusion clustering [44], which partitions datasets into clusters such that no two cluster centers are within a specified distance of each other. Detailed methodologies of these splitting strategies are elaborated in section 5 of the Supplementary Information.

For the evaluation of the interpretability on protein sequences, we identified 1,125 experimental drug-protein complex structures from the PDB (Protein Data Bank, https://www.rcsb.org/) network, which are included in the BindingDB dataset. Then, 1,862 drug-protein pairs corresponding to these experimental complex structures are set aside as the test set, while the remaining pairs are randomly split into the training and validation sets.

Typically, binary classification problems employ AU-ROC (Area Under the Receiver Operating Characteristic Curve) and AUPRC (Area Under the Precision-Recall Curve) as gold standard metrics for model performance evaluation. Additionally, accuracy, sensitivity, and specificity at the threshold of the best F1 score are also used for comparison purposes. Moreover, we introduce the Top-N hit metric to evaluate the DTI model’s capability to identify actual interactive protein fragments. By defining amino acids within 6 Å of the nearest ligand molecule as protein pockets, this metric gauges the likelihood of hitting the protein pocket at least once among the top N (where N could be 1, 3, 5, 10) attentive amino acids in each protein sample. A higher Top-N hit implies that the DTI model is more adept at learning the actual interactive protein fragments in the given sample.

In the end, we conduct five independent runs with different random seeds to ensure stability and robustness. Five-fold cross-validation results and relevant statistical results are also provided in Supplementary Information.

## 3 METHODS

### 3.1 Data preparation

We advocate the combination of protein embedding from a deep LLM and a drug embedding through shallower networks, thereby establishing an unbalanced dual-branch network. The rationale behind this design lies in the contrasting capabilities of deep and shallow networks. Given that amino acids bound in topological space may not align sequentially, deep LLM offers a broad receptive field capable of encompassing multiple amino acids constituting local spatial substructures. In contrast, the graph structure of a drug molecule explicitly delineates spatial properties and crucial functional groups could be covered by a relatively small receptive field. Employing shallow networks could prevent overfitting. Illustratively, in Fig.1d, the inadequacy of a 3-layer network in extracting topologically bounded amino acids in Acetylcholinesterase is evident despite its success in capturing the pyridine-carbonyl (the interactive functional group in complex 1GPK) of Huperzine A drug. Conversely, a 24-layer network (usually the average depth of LLMs) effectively focuses on the spatial properties of proteins but obscures critical nodes of drug molecules (Fig.1e).

To generate protein target features, we utilize a pretrained protein LLM to produce embeddings 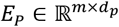, where *m* is the length of a protein and *d*_*p*_is the preset dimension of the hidden layer. ESM [37] and ProtBert [38] (separately used in distinct downstream tasks) are selected to process the original sequence data. Our input drug molecules are represented using SMILES. We preprocess SMILES into graphs incorporating node features and adjacency matrices using RDkit. Each node’s feature is expressed through one-hot encoding, encompassing the atom type, degree, formal charge, number of radical electrons, hybridization, aromaticity, and the total hydrogen count for the atom. Consequently, this processing yields a drug ligand embedding, 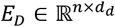, where *n* is the length of the drug and *d*_*d*_is 74. During the training process, the maximum lengths of protein and drug are set to 1200 and 290, respectively. Additionally, we explored the utilization of other protein and drug LLMs; all ablation experiments on both protein and drug LLMs can be found in the Supplementary Information.

### 3.2 Feature extractor

GCNs are potent neural architectures explicitly tailored to process graphs, leveraging their structural information effectively. For the drug compound, we fed the molecular graph *E*_*D*_into a three-layer GCN module to capture the information on non-hydrogen heavy atoms and functional groups. The message propagation mechanism of GCN ensures weighted aggregation of information concerning topologically bonded atoms (chemical bonds). The deeper layers employ superposition to broaden the receptive field, accurately extracting and characterizing substructures within small drug molecules. The formulation of the GCN extractor utilizes a convolution operation described as follows:

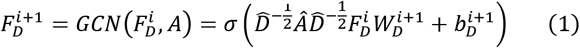

Where *A ∈ ℝ*^*n×n*^ is the adjacency matrix of the drug graph (*n*is the number of the nodes in the graph), *I*is the identity matrix, and 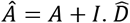 is the diagonal node degree matrix calculated from A. 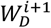 and 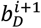 are the earnable weight matrix and bias vector of the layer i+1, respec-tively, and the output is 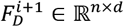. It should be noted that 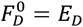, the initial embedding vector of the drug.

Besides, we apply a linear layer to adjust the dimension of protein embedding *E*_*p*_to *F*_*p*_*∈ ℝ*^*m×d*^.

### 3.3 Attentive aggregation

The attention mechanism, well-known in Transformer architectures [45] for machine translation, has raised considerable research interest. The mechanism typically comprises three vectors, query vector (q), key vector (k), and value vector (v), to compute interaction values (e.g., dot products or correlation coefficients) between the query and key vectors to weight the summation of the value vectors [46-48]. A classic multi-head attention mechanism can be represented by eq.2-4.

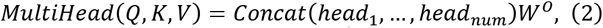

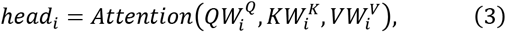

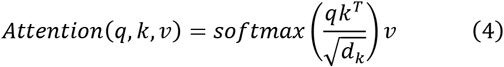

Where *d*_*k*_is the dimension of vectors q and k, num is the preset number of attention heads, 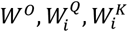, and 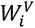 are trainable parameters.

We have designed a novel attentive aggregation architecture (shown in Fig.1b) specifically aimed at capturing the pairwise interactions between protein and drug substructures. As the feature representations of proteins *F*_*p*_and drugs *F*_*D*_share the same dimension *d*, they could be viewed as sentences (token sets) of varying lengths. We first randomly initialize a token *h ∈ ℝ*^1*×d*^ as the head of *F*_*p*_and *F*_*D*_, respectively, and then define 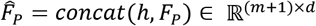 and 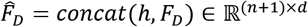. In the two parallel branches, 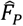 and 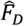 each is calculated by a distinct 1-layer Transformer. The updated heads are retrieved to serve as the query vectors for calculating cross-attention in the opposite branch. Outputs from four positions are summed with pre-set weights to the original token *h*. We retain the original head *h*to ensure the smoothness of model predictions. The mathematical representation of this process is as follows:

Transformer step:

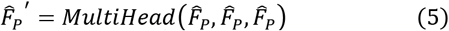

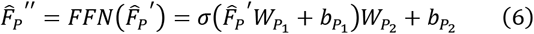

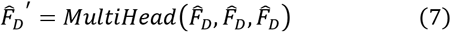

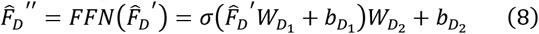

Where *W*_*p_*1_, *W*_*p_*2_, *b*_*p_*1_, *b*_*p_*2_, *W*_*D_*1_, *W*_*D_*2_, *b*_*D_*1_, and *W*_*D_*2_are trainable parameters.

**Cross-attention step:**

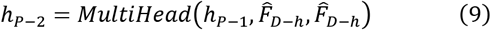

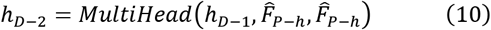

Where *h*_*p*−1_and 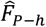 are the head token and rest tokens of 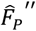, respectively, and *h* and 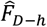 are the head token nd rest tokens of 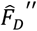 respectively. Thus we could calculate the final output _*p*_^0^.

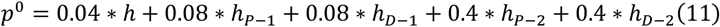

### 3.4 Decoder and loss function

For the calculated *out*_*p*_*ut*_*token*_*∈ ℝ*^1*×d*^, we fed it into a 3-layer multilayer perceptron to compute the interaction probability. The calculation process in each layer can be represented by the following formula.

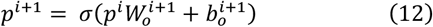

Where the predictive score _*p*_ *= sigmoid*(_*p*_^3^).

Finally, the cross-entropy loss function is used to minimize the distance between labels and predictions.

### 3.5 Cross-domain module

As shown in Fig.1c, the MCD module introduces an additional decoder to implement adversarial learning while maintaining the core architecture. MCD achieves domain alignment from the target to source domains at a low computational cost, minimizing the impact on model interpretability.

Given a source domain 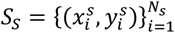 of *N* labeled drug–target pairs and a target domain of 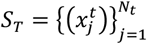 of *N*_*t*_unlabeled drug–target pairs, UdanDTI could be decomposed as two decoders with different initialization methods, *F*_1_ and *F*_2_, and a main architecture *G*. MCD module repeats the following three steps iteratively until convergence. In Step 1, *G F*_1_ and *F*_2_ are trained simultaneously on *S*_*S*_to ensure the classification accuracy of the model in the source domain. In Step 2, *G* remains fixed while two diverse classifiers are trained on *S*_*T*_to widen the discrepancy. Notably, Step 1 is simultaneously included to restrict the direction of the decoder hyperplane changes. In Step 3, parameters in classifiers are kept constant and *G* is trained on *S*_*T*_to reduce divergence (samples with low confidence) between them by extracting improved features. Through this alternating process of divergence and compromise, UdanDTI_MCD_ enables *G* to learn domain-invariant features that are robust to different decision boundaries. Finally, we choose the converged model obtained after 200 epochs.

## 4 RESULTS

### 4.1 In-domain experiment

In the in-domain experimental setup, we conduct a comparative analysis against seven state-of-the-art models. We obtained results for Random Forest (RF), DeepConv-DTI, GraphDTA, MolTrans, and DrugBAN from cited references [15]. Additionally, MGraphDTA and MCANet were replicated and deployed utilizing publicly available codes.

Table 1 summarizes the comparative results on the Bind-ingDB and BioSNAP datasets. Across all evaluation met-rics—AUROC, AUPRC, accuracy, sensitivity, and specificity—UdanDTI consistently demonstrates superior performance over the baseline models. These results suggest that UdanDTI captures crucial information from fused dual-branch features more effectively than learning from data or descriptors based on statistical learning. We also designed ablation experiments on BioSNAP to evaluate the individual contributions of the unbalanced dual-branch and attentive aggregation module (which can be found in the Supplementary Information).

**Table 1.** In-domain comparison between UdanDTI and seven other advanced DTI models on the BindingDB and BioSNAP datasets.

| Dataset | Method | AUROC | AUPRC | Accuracy | Sensitivity | Specificity |
| --- | --- | --- | --- | --- | --- | --- |
| BindingDB | RF | 0.942±0.011 | 0.921±0.016 | 0.880±0.012 | 0.875±0.023 | 0.892±0.020 |
|  | DeepConv-DTI | 0.945±0.002 | 0.925±0.005 | 0.882±0.007 | 0.873±0.018 | 0.894±0.009 |
|  | GraphDTA | 0.951±0.002 | 0.934±0.002 | 0.888±0.005 | 0.882±0.012 | 0.897±0.008 |
|  | MolTrans | 0.952±0.002 | 0.936±0.001 | 0.887±0.006 | 0.877±0.016 | 0.902±0.009 |
|  | MGraphDTA | 0.951±0.002 | 0.936±0.002 | 0.888±0.006 | 0.881±0.017 | 0.899±0.011 |
|  | MCANet | 0.956±0.001 | 0.940±0.003 | 0.892±0.003 | 0.887±0.011 | 0.904±0.004 |
|  | DrugBAN | 0.960±0.001 | 0.948±0.002 | 0.904±0.004 | 0.900±0.008 | **0.908±0.004** |
|  | UdanDTI | **0.965±0.001** | **0.955±0.001** | **0.911±0.004** | **0.920±0.007** | 0.900±0.007 |
| BioSNAP | RF | 0.860±0.005 | 0.886±0.005 | 0.804±0.005 | 0.823±0.032 | 0.786±0.025 |
|  | DeepConv-DTI | 0.886±0.006 | 0.890±0.006 | 0.805±0.009 | 0.760±0.029 | 0.851±0.013 |
|  | GraphDTA | 0.887±0.008 | 0.890±0.007 | 0.800±0.007 | 0.745±0.032 | 0.854±0.025 |
|  | MolTrans | 0.895±0.004 | 0.897±0.005 | 0.825±0.010 | 0.818±0.031 | 0.831±0.013 |
|  | MGraphDTA | 0.896±0.006 | 0.899±0.004 | 0.811±0.012 | 0.785±0.035 | 0.856±0.018 |
|  | MCANet | 0.901±0.006 | 0.899±0.005 | 0.827±0.009 | 0.810±0.035 | 0.827±0.011 |
|  | DrugBAN | 0.903±0.005 | 0.902±0.004 | 0.834±0.008 | 0.820±0.021 | 0.847±0.010 |
|  | UdanDTI | **0.941±0.003** | **0.942±0.004** | **0.876±0.006** | **0.876±0.016** | **0.877±0.009** |

The results on the Human dataset, depicted in Fig.2a and 2b, indicate that most models show promising performance under random splitting, both in AUROC and AUPRC. However, a previous study [26] has highlighted the risk of overfitting in the Human dataset. To mitigate this risk, we adopted a cold-pair splitting strategy for further evaluation. Fig.2a and 2b illustrate a notable decline in the AUROC and AUPRC performance of all models when transitioning from random segmentation to cold-pair splitting. Under this strategy, UdanDTI demonstrates a clearer competitive edge than the other models, consolidating its performance advantage.

**Fig 2.**
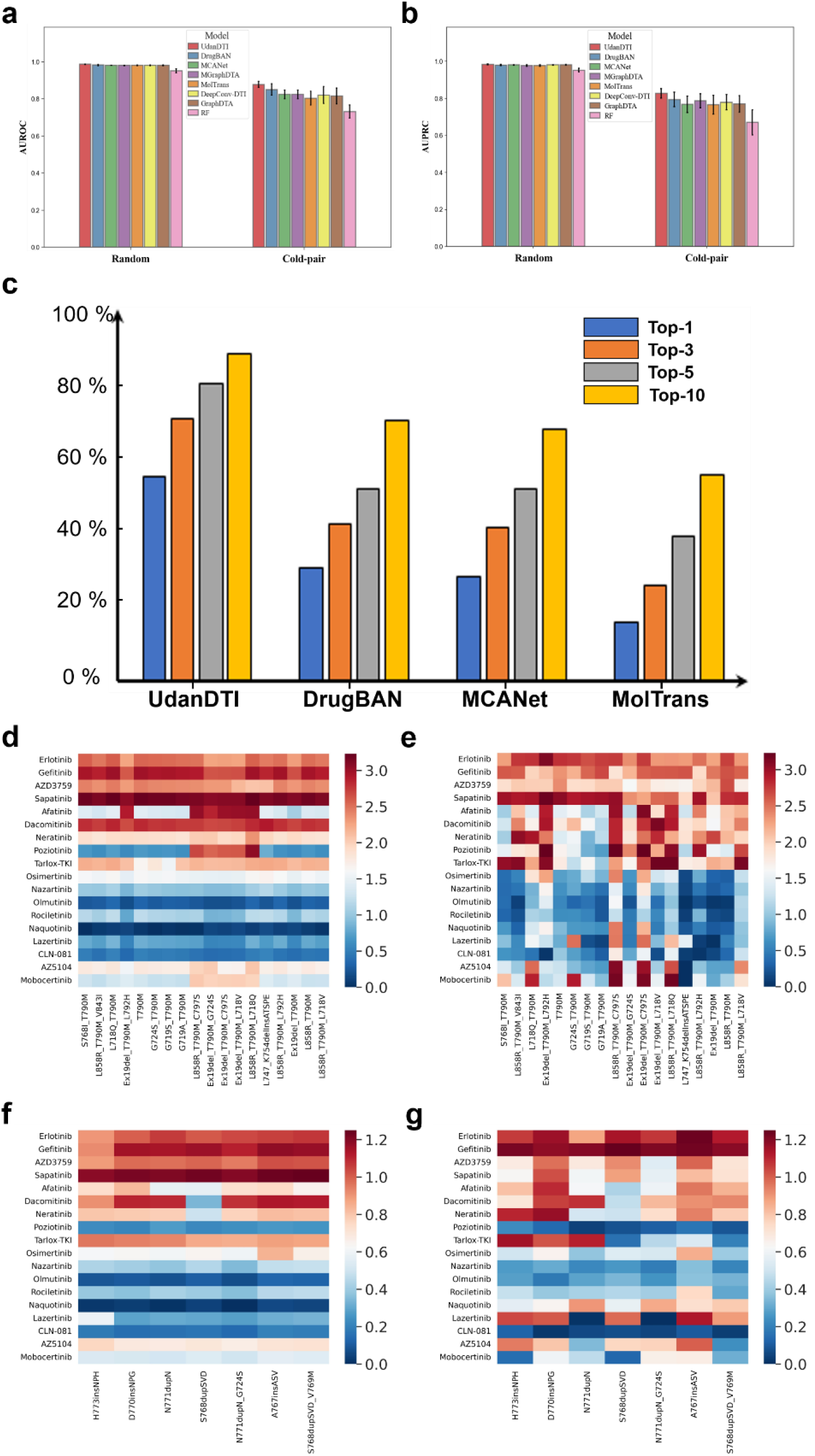
(a) and (b) indicate the comparison between UdanDTI and seven other advanced DTI models on the Human dataset. (a) AUROC performance under random splitting and cold-pair splitting. (b) AUPRC performance under random splitting and cold-pair splitting. (c) Performance comparison between UdanDTI and other interpretable models under the Top-N hit metric. (d-g) show the drug-protein response values predicted by UdanDTI and measured by in-vitro experiments., including predicted results among T790M-like subgroup (d) and Ex20ins-L subgroup (f), and in-vitro experimental results [35] among T790M-like subgroup (e) and Ex20ins-L subgroup (g).

### 4.2 Cross-Domain Experiment

The in-domain setting may not authentically represent the true performance of DTI predictive models in prospective forecasting. As previously highlighted, a drug-bias trap in the DTI dataset may lead to accurate predictions solely based on drug characteristics, overlooking actual interaction patterns. The apparent high accuracy might stem from biases toward drugs and overfitting grounded in protein homology rather than a model’s genuine predictive performance.

In real-world scenarios, the vast chemical space often contains drug-target pairs that are Out-of-Distribution (OOD) with the training data. Employing the same source and target domain setup as in DrugBAN, latent features of proteins and drugs in the test set are explicitly distanced from the feature distribution in the training set through clustering methods. To confront the cross-domain challenges, we activated the MCD module of UdanDTI. Table 2 presents the performance evaluation of the BindingDB and BioSNAP datasets under cluster-based pair segmentation. All DTI models showed a notable decrease in performance due to the minimized information overlap between the training and test sets. Under the cross-domain setting, vanilla UdanDTI demonstrated robustness, exhibiting AU-ROC values 6% and 10% higher than DrugBAN on the BioSNAP and BindingDB datasets, respectively. Random Forest (RF) exhibited commendable performance in the cross-domain settings, surpassing some deep learning baselines (DeepConv, GraphDTA, and MolTrans), likely due to its reliance on statistics-based fingerprint features. Meanwhile, deep learning models emphasizing interactive learning (MCANet, DrugBAN, and UdanDTI) exhibited superior predictive performance. Further refinement and tuning of the MCD module improved UdanDTI’s generalization capabilities, enhancing AUROC by 8% and 5% on the two datasets, respectively. These outcomes highlight UdanDTI’s capability to address real-world challenges. Additionally, we provide an ablation experiment of various cross-domain modules and the attentive aggregation module in the Supplementary Information.

**Table 2.**
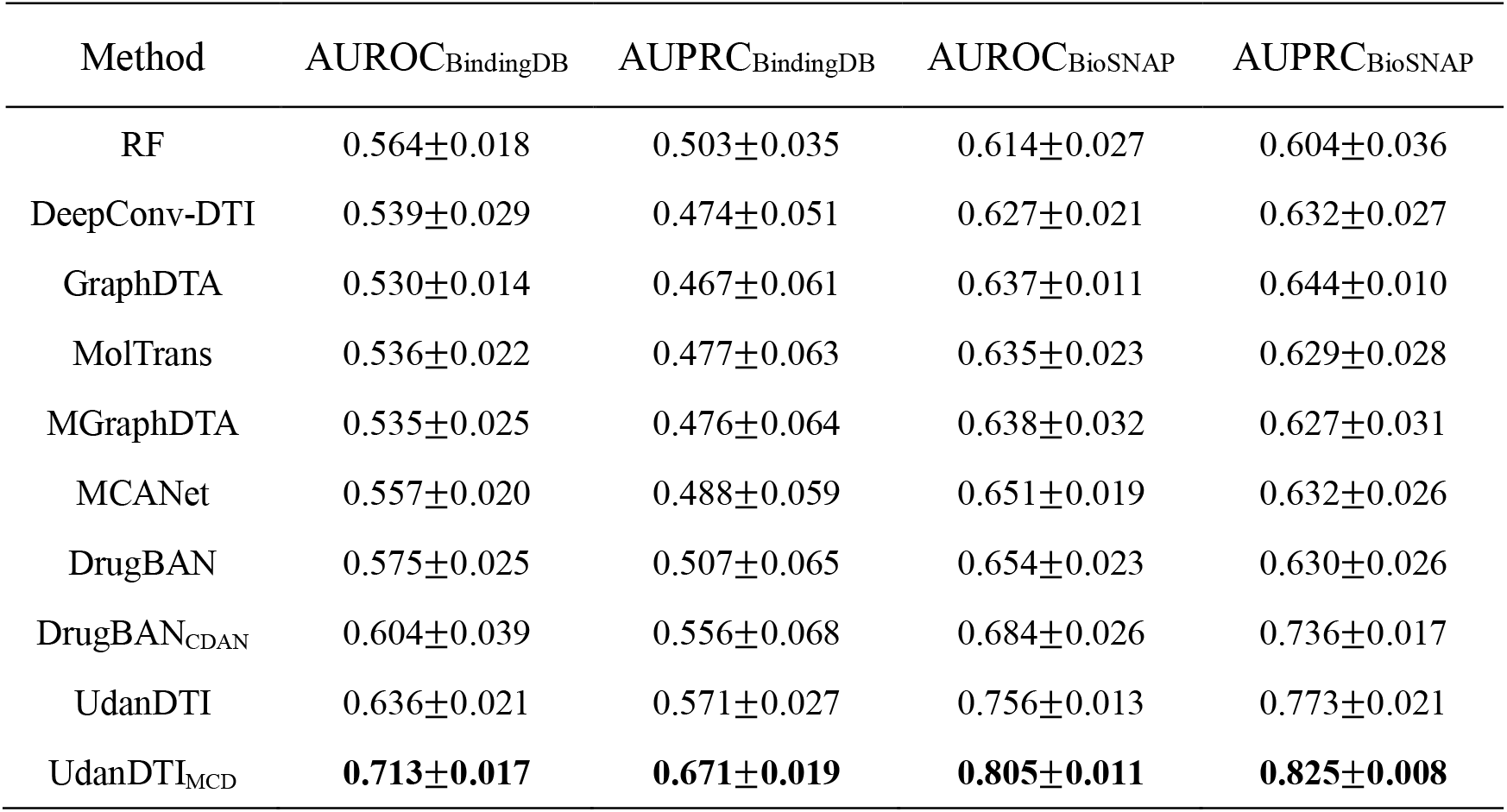
Cross-domain comparison between UdanDTI and eight other advanced DTI models on the BindingDB and BioSNAP datasets.

| Method | AUROC<sub>BindingDB</sub> | AUPRC<sub>BindingDB</sub> | AUROC<sub>BioSNAP</sub> | AUPRC<sub>BioSNAP</sub> |
| --- | --- | --- | --- | --- |
| RF | 0.564±0.018 | 0.503±0.035 | 0.614±0.027 | 0.604±0.036 |
| DeepConv-DTI | 0.539±0.029 | 0.474±0.051 | 0.627±0.021 | 0.632±0.027 |
| GraphDTA | 0.530±0.014 | 0.467±0.061 | 0.637±0.011 | 0.644±0.010 |
| MolTrans | 0.536±0.022 | 0.477±0.063 | 0.635±0.023 | 0.629±0.028 |
| MGraphDTA | 0.535±0.025 | 0.476±0.064 | 0.638±0.032 | 0.627±0.031 |
| MCANet | 0.557±0.020 | 0.488±0.059 | 0.651±0.019 | 0.632±0.026 |
| DrugBAN | 0.575±0.025 | 0.507±0.065 | 0.654±0.023 | 0.630±0.026 |
| DrugBAN<sub>CDAN</sub> | 0.604±0.039 | 0.556±0.068 | 0.684±0.026 | 0.736±0.017 |
| UdanDTI | 0.636±0.021 | 0.571±0.027 | 0.756±0.013 | 0.773±0.021 |
| UdanDTI<sub>MCD</sub> | **0.713±0.017** | **0.671±0.019** | **0.805±0.011** | **0.825±0.008** |

### 4.3 Quantitative Comparison of Interpretability on Protein

Addressing branch bias is a common challenge in multibranch learning frameworks. While some black-box models directly concatenate protein and drug representations, achieving high accuracy, their practical significance is relatively limited. The pervasive drug bias trap has emerged as a substantial obstacle for ML-based DTI models [22, 34]. We highlighted that it is partly a trap caused by the dual-branch network paradigm and the relatively simple struc-ture of drug molecules

To evaluate the efficiency of models in learning interaction patterns, we utilized a Top-N hit metric to assess model performance. Although some other methods [49, 50] have been used to measure the interpretability, we choose a more intuitive quantitative comparison: the overlap between the attentive scores predicted by the model and the actual domains of protein pockets. We reproduced models such as MCANet, MolTrans, and DrugBAN based on their respective paper details, trained them on the BindingDB training set, and then evaluated their Top-N hit on 1,862 DTI pairs with actual structures. The experimental results, displayed in Fig.2c, unequivocally demonstrate UdanDTI’s superior performance compared to other advanced models. We further compared the frequency of effective hits of different amino acids. The experimental results (found in Supplementary Information) show that UdanDTI is more effective in predicting the interactions between heavy atoms on the side chains of PHE, LEU, MET, and HIS amino acids and ligands. The specially designed attentive aggregation module facilitates an intuitive exploration of the influence exerted by the protein and drug substructures on the predicted results. By solely applying Large Language Models (LLM) to the protein branch, UdanDTI effectively balances both branches. These results suggest that UdanDTI relatively mitigates the drug-bias trap, and its interpretability is substantiated through quantitative experiments

### 4.4 Prediction on T790M and EX20 subgroup

EGFR mutations commonly occur in exon 18-21 and are primary driver mutations identified in non-small cell lung cancer (NSCLC). Researchers categorized EGFR mutants into four subgroups based on sensitivity and structural changes [43]. Predicting interactions between protein mutants and drugs presents an immensely challenging task due to the tendency of EGFR gene mutations to occur predominantly on a single exon or even a single amino acid. Moreover, the public benchmark DTI datasets contain solely the wild EGFR gene and a limited set of early-developed drugs (Afatinib, Erlotinib, Gefitinib, and Osimertinib), further complicating predictions. To facilitate domain transfer learning, we converted the experimental mutant-drug affinities to binary values (0-1) and activated the MCD module, also comparing the performance with DrugBANCDAN.

Results indicate that in T790M-like and Ex20ins-L subgroups, where gene mutations significantly modify the protein-binding pocket’s structure, resulting in resistance, UdanDTI demonstrated robust performance. UdanDTI achieved an AUROC of 0.83 and an AUPRC of 0.91 in the T790M-like subgroup, outperforming DrugBANCDAN by 7% and 4%, respectively. In the Ex20ins-L subgroup, UdanDTI attained an AUROC of 0.79 and an AUPRC of 0.86, surpassing DrugBANCDAN by 8% and 9%, respectively.

Fig.2d-2g presents a visualization comparison between UdanDTI’s predictions and the experimental results in the original article [43]. To facilitate comparison, we sorted the response and predicted values of all protein-drug pairs and color-coded the corresponding cells accordingly. Visualization results show that the effect of the drug-bias trap was further reinforced due to the concealment of genetic mutations. For pairs of the same drug and different mutants, DTI models often result in approximate predictions. Notably, UdanDTI successfully predicted a series of exon-20 missing proteins in the T790M subgroup and a series of S768dup proteins in the Ex20ins-L subgroup, partially escaping the amplified drug bias and achieving accurate predictions. However, challenges persisted in accurately predicting the interactions with drugs Naquotinib, Lazertinib, and AZ5104, primarily due to their complex structures that are located far from the training set in the feature space. This underscores the need for further enhancements in model designing and feature representation to better capture these complex interactions.

### 4.5 Interpretability with Attention Visualization

UdanDTI, equipped with an attentive aggregation module, explicitly learns the molecular substructure contributions of both proteins and drugs to the final predictions. Visualizing the attention weights provides insight into the molecular and amino acid levels. We categorized the cocrystalized ligands into three groups for analysis: Classical, Discrete fragments, and Complex structure, and visualized two top predictions for each category.

Classical samples typically contain proteins of moderate length and drugs with simple structures, with fewer binding points and pockets relatively loose. For PDB structure 2B17 (DIF compound bound to Phospholipase A2 protein), UdanDTI correctly identified the carboxyl group and 22th amino acid TYR involved in the binding patterns, and TYR also forms hydrogen bonds with secondary amines, which is considered as an important amino acid. The 30th GLY, which is involved in the arene-H reaction, is also considered important. Unfortunately, the 49th ASP, which reacts with carboxyl and two water molecules, has not been correctly identified. For PDB structure 3FY8 (XCF compound bound to Dihydrofolate reductase), UdanDTI accurately identified the amino group and unilateral carbon atoms involved in the reaction at the 2, 4-diaminopyrimidine site. UdanDTI recognizes the amino acids, 5th LEU and 92th PHE, that react with an amino group on one side and the amino acid binding point at the 22nd TRP, which binds to carbon atoms. However, it ignores the 30th, 109th, and 111th amino acids that react with an amino group on the other side.

Discrete fragments samples refer to pockets of a target protein often composed of non-sequentially adjacent amino acid fragments. This category challenges DTI models to discern the 3D spatial structure from 1D protein sequence data. We examine the results of UdanDTI in two highconfidence samples in the following. In the PDB structure 3G5K (a complex of Actinonin drug and peptide deformylases), UdanDTI correctly predicted almost all interaction patterns. The protein fragments that form the binding site (51th Val 57th Gln, 85th Arg 89th Pro, and 110th Phe 115th Glu), and the carbonyl and hydroxyl groups involved in the binding reaction are all highlighted in red. Unfortunately, UdanDTI ignored the reaction of the 157th Glu with the amyl carbon atom, possibly because the attention mechanism amplifies those more significant interactions. Similarly, in the PDB structure 2DRC (a complex of Methotrexate drug and Dihydrofolate Reductase), UdanDTI identified multiple protein fragments surrounding the ligand molecule (5th Ile - 9th Ala, 19th Ala - 27th Asp, 44th Arg - 48th Glu, 52th Arg - 57th Arg, 93th Val 97th Gly, 117th Ala 121th Gly). It indicated that UdanDTI identified potential pockets based on drug molecules. For specific binding patterns, UdanDTI accurately identified the 52th Arg and 57th Arg as sidechain donors of benzoyl and glutaric acid, and the 5th Ile and 94th Ile as the backbone acceptors of 2, 4-diaminopteridine. However, the model partially identified the triangular interactions of 27th Asp, 2, 4-diaminopteridine, and 113th Thr, exposing the inadequacy of the cross-attention mechanism in the complex response.

Complex structure samples involve target proteins with intricate spatial configurations that may confound DTI models. As shown in Fig.3, only one reaction between the 276th Leu and the benzoic acid is observed in the PDB structure 4P38 (complex of 21T and Corticosteroid 11-beta-dehydrogenase isozyme 1). UdanDTI highlighted multiple amino acids neighboring ligands, including 276Leu and the carboxyl carbon atom of benzoic acid. In the PDB structure 8F4W (a complex of VFX and N-acetyltransferase Eis), the ligand reacts primarily with amino acids on the 26th Asp 36th Trp protein fragments, which UdanDTI accurately captures. However, UdanDTI performs relatively poorer in the ligand molecules. It only extracts the binding patterns of the alcohol groups on dimethylamino and cyclohexane with their corresponding amino acids, missing the important influence of methoxyphenyl groups in the interaction. Such polycyclic and benzene structures challenge the ability of GNN.

**Fig 3.**
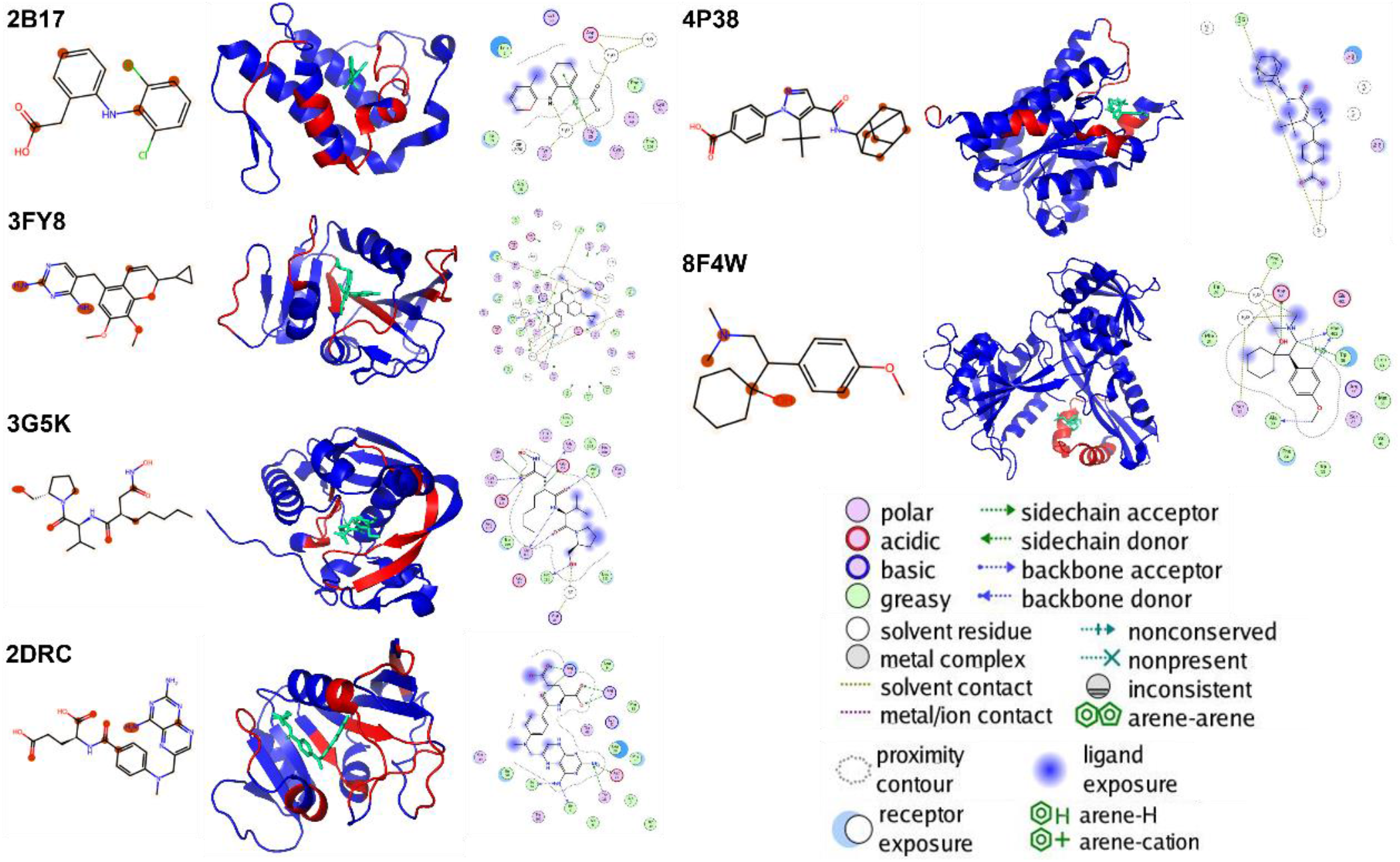
Visualization results of UdanDTI. For each sample, we visualize the ligand molecule using RDKit and color the top 20% of the atoms that UdanDTI emphasizes in red (left sub-figure). Using PyMol, we illustrate the actual co-crystal structure obtained from the PDB library and highlight the top ten weighted amino acids with their preceding and following amino acids, all in red (middle sub-figure). Finally, we utilize Molecular Operating Environment (MOE) software to visualize the actual ligand molecular reaction diagram as a comparative standard (right sub-figure).

### 4.6 Complementary Tool for Docking

Molecular docking is a computational technique used to predict the preferred binding poses of ligands (drugs) to receptors (proteins) under specific conditions. DifDock [51] Innovatively adopts a diffusion model to simulate the uncertainty in molecular binding in real scenarios. It introduces a noise process by sampling positional changes, conformational flips, and bond twists (steps, turns, and twists) in ligand conformation and then trains a denoising model by feeding the altered ligand conformation and protein conformation.

In our experiments, we utilized the pre-trained DiffDock model on 1,125 drug-protein pairs with experimentally validated complex structures (detailed in 3.2) to infer the most probable ligand poses (rank_1 provided by confidence predictor of DiffDock). We further refined this approach by restricting the input to DiffDock to only include amino acids covered by the top 20% of UdanDTI’s predictions. This focused input allowed DiffDock to achieve higher accuracy in ligand pose prediction, with the average RMSD improving from 6.55 Å to 5.24 Å.

As shown in Fig.4, we chose two representative examples to elucidate. In complex 5YVC (Structure of CaMKK2 in complex with CKI-012, as shown in Fig.4a), a narrow but precisely targeted perspective assisted DiffDock in better focusing on the protein pocket, resulting in more precise ligand pose generation. Excluding the terminal propionic acid, the ligand generated by DiffDock with UdanDTI’s assistance exhibited better alignment with the native small molecule. In complex 7X9D (DNMT3B in complex with harmine, as shown in Fig.4b), the amino acid range provided by UdanDTI effectively avoided other potential misleading protein pockets. While the ligand pose was not pixel-perfect, it was accurately positioned within the correct pocket, illustrating UdanDTI’s ability to mitigate some of DiffDock’s limitations in positional changes.

These results indicate that UdanDTI can supplement docking models like DiffDock in molecular docking tasks. Furthermore, we include comparison experiments with DiffDock on time performance and virtual screening capability (assessed manually) in the Supplementary Information section.

**Fig 4.**
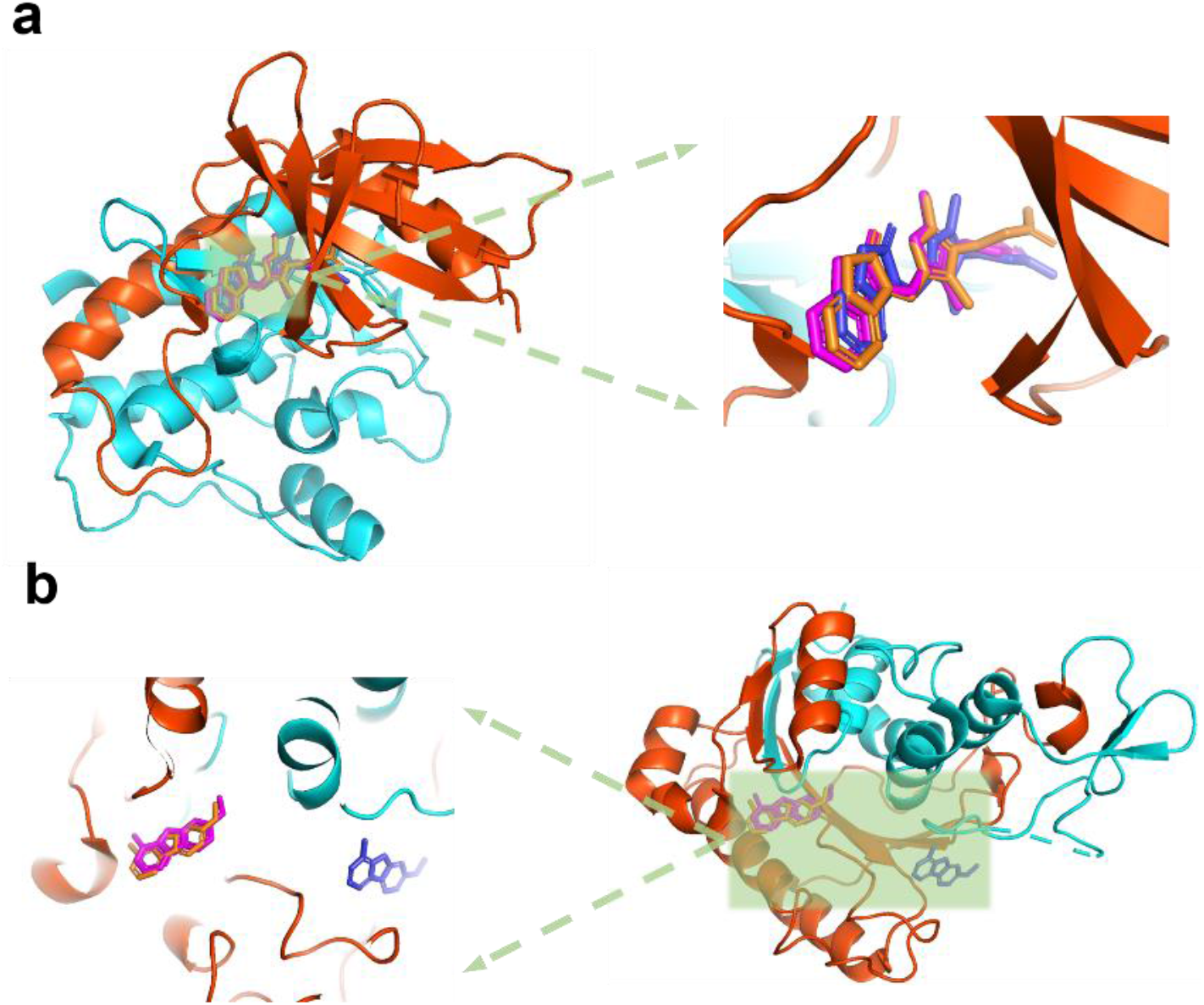
Two representative examples of UdanDTI-assisted DiffDock docking. The range of amino acids selected by UdanDTI on the protein is highlighted in orange. The ligands inferred by UdanDTI-assisted DiffDock, directly predicted by DiffDock, and experimentally measured were colored in orange, blue, and magenta, respectively. (a) UdanDTI assists DiffDock in inferring a more precise binding pose. (b) UdanDTI assists DiffDock in avoiding other erroneous protein binding pockets.

## 5 CONCLUSION

This paper introduces UdanDTI, an advanced deep learning architecture designed for predicting DTIs. Through comprehensive experiments, UdanDTI has demonstrated exceptional performance on diverse public datasets, outperforming existing state-of-the-art DTI models across both in-domain and cross-domain evaluations. Furthermore, UdanDTI serves as a crucial auxiliary tool for molecular docking, enhancing the accuracy of ligand docking by identifying active protein fragments.

A significant aspect of our study is the identification and mitigation of drug-bias trap caused by balanced dual-branch architectures. This trap leads to over-reliance on the drug branch, hindering the interpretability and widespread application of existing DTI models. UdanDTI’s distinctive unbalanced dual-branch and attentive aggregation module mitigates the drug-bias trap, providing advanced interpretability from a biological standpoint. This architecture allows for the visualization of molecular interaction patterns, offering valuable insights into drug-protein interactions at both atomic and amino acid levels. Our work represents the first quantitative comparison of DTI model interpretability on proteins, demonstrating UdanDTI’s ability to capture the locally active protein fragments and potential protein pockets.

The introduction of cross-domain module and LLM significantly enhances UdanDTI’s generalization ability. UdanDTI_MCD_ achieved 11% and 12% higher AUROC than the best DTI model on two public datasets, BindingDB and BioSNAP, under cross-domain setting, respectively. Additionally, UdanDTI excels in predicting drug responses among two crucial subgroups of EGFR mutants. The flexibility of the UdanDTI framework suggests its potential adaptability to a wide range of biological interaction challenges, including protein-protein interactions.

While our study did not incorporate highly accurate 3D structural protein data, given that only a few known protein sequences have detailed structural information. Recent advancements in protein 3D structure prediction, like AlphaFold [52], show promising strides. However, direct integration of predictive structures into the DTI framework may present challenges. Given the confidence score provided by AlphaFold, a composite framework considering the availability and reliability of predictive structures might offer a more viable solution to address the DTI problem effectively.

## ACKNOWLEDGMENT

This work is supported by the National Key R&D Program of China (2021YFF1201303, 2022YFC2703105), Guoqiang Institute of Tsinghua University, and Beijing National Research Center for Information Science and Technology (BNRist). The funders had no roles in study design, data collection and analysis, the decision to publish, and manuscript preparation.

---

Pei-dong Zhang received the BSc degree in automation from Shanghai Jiao Tong University, China, in 2020, and the MSc degree in electronic information from Shanghai Jiao Tong University, in 2023. Currently, he is working toward the PhD degree in computer science and technology at Tsinghua University. His research interests include computer-aided drug discovery and deep learning.

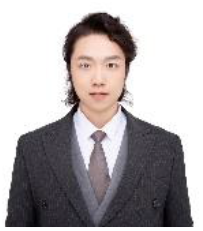

Jianzhu Ma is currently an associate professor at the Institute for AI Industry Research, Tsinghua University, and a recipient of the Overseas Young Talent Program. He received his Ph.D. from Toyota Technological Institute at Chicago in 2016, followed by a position as a project scientist at the University of California, San Diego. From 2020 to 2021, he served as a Walther Assistant Professor in the Department of Computer Science and Biochemistry at Purdue University, and later as an associate professor at Peking University’s AI Institute and School of Public Health. His research focuses on artificial intelligence, systems biology, biopharmaceuticals, and smart healthcare. He has received multiple awards, including the Best Paper Award at RECOMB, the Warren DeLano Award at ISMB, and Best Poster Award at the RNA and Protein Folding Conference. He has published in top journals like Nature Methods and Nature Machine Intelligence, with some of his work featured as cover articles.

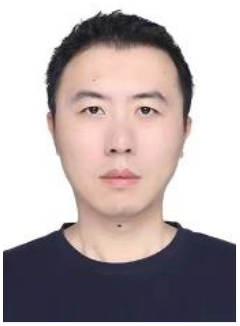

Ting Chen biography is a professor in the Department of Computer Science and Technology at Tsinghua University. He earned his Bachelor’s degree from Tsinghua University in 1993 and a Ph.D. from Stony Brook University in 1997. He was a lecturer at Harvard Medical School and later held faculty positions at the University of Southern California, where he directed the Computational Biology Division. His research focuses on big data algorithms and machine learning in genomics and proteomics. He has published over 120 papers in top journals like Cell, Science, and Nature Communications, with over 10,000 citations, and has led projects funded by NSF, NIH, and NSFC. He received the Sloan Research Fellowship in 2004. appears here.

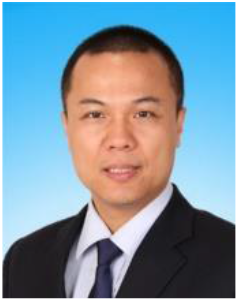

## Supplementary Information

## 1 Dataset statistics, training procedure, and hyperparameter setting

Table S1 presents the statistical features of the experimental datasets. Table S2 shows the training procedure, with the notations and explanations. Table S3 details the hyperparameters utilized in UdanDTI.

**Table S1.**
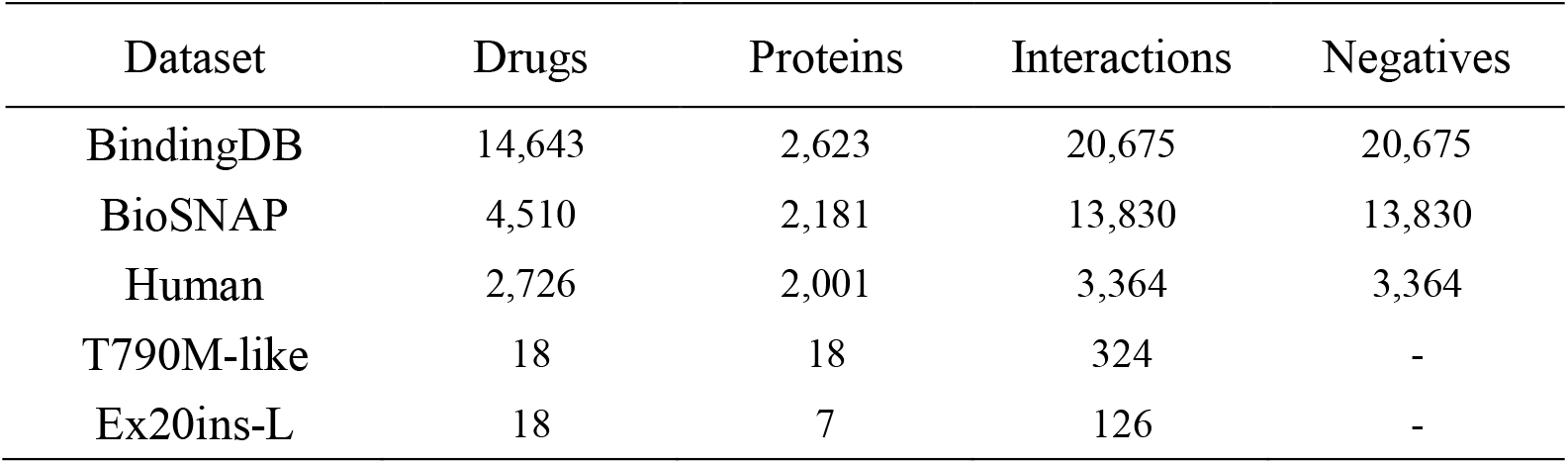
Datasets statistics.

| Dataset | Drugs | Proteins | Interactions | Negatives |
| --- | --- | --- | --- | --- |
| BindingDB | 14,643 | 2,623 | 20,675 | 20,675 |
| BioSNAP | 4,510 | 2,181 | 13,830 | 13,830 |
| Human | 2,726 | 2,001 | 3,364 | 3,364 |
| T790M-like | 18 | 18 | 324 | - |
| Ex20ins-L | 18 | 7 | 126 | - |

**Table S2.**
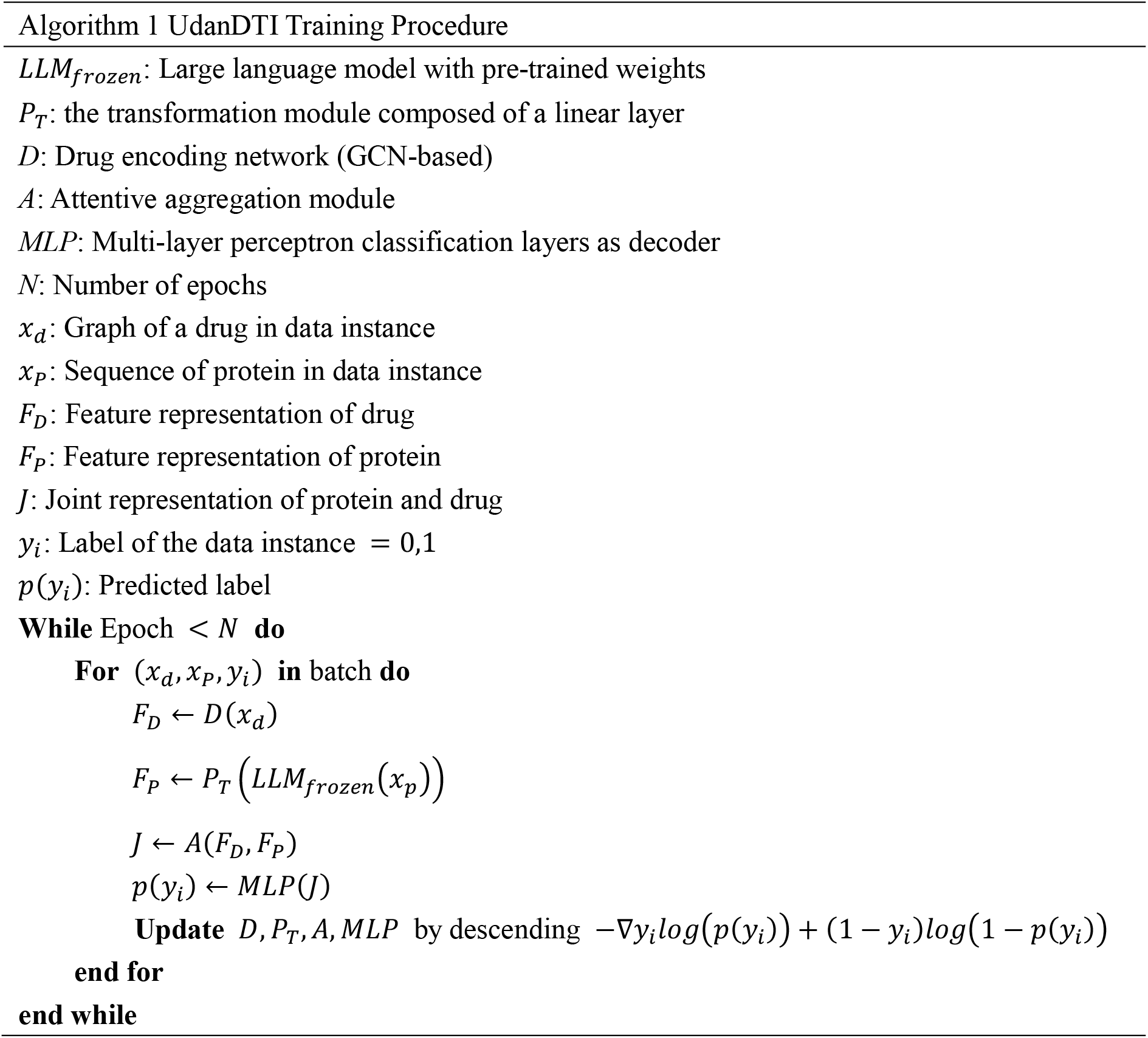
Training procedure and notations.

For direct comparative analyses between UdanDTI and the state-of-the-art (SOTA) machine learning-based methods, we employed three publicly available drug-target interaction (DTI) datasets: BindingDB, BioSNAP, and Human. BindingDB[1], an expansive open database, primarily focuses on interactions between proteins identified as potential drug targets and small drug-like ligands, offering a broader scope than the other databases. In our experiments, we utilized a preprocessed, low-bias subset of the BindingDB dataset[2], comprising 14,643 drugs and 2,623 proteins. The BioSNAP[3] dataset encapsulates information on genes targeted by drugs available on the US market. Derived from the DrugBank database and balanced by Huang et al.[4], this dataset includes 4,510 drugs and 2,181 proteins. The Human dataset crafted by Liu et al.[5] integrates highly reliable negative samples of compound-protein pairs obtained through a systematic screening framework. We employed a balanced version consisting of 2,726 drugs and 2,001 proteins, which is consistent with prior studies and ensures an equal number of positive and negative samples.

Additionally, researchers compiled mutations in 16,715 non-small cell lung cancer (NSCLC) patients with EGFR mutations, computed the interaction spectra of 76 EGFR protein mutants with 18 drugs, and further categorized these mutants into four subgroups[6]: 1. Classical-like mutations that are distant from the ATP-binding pocket (Classical-like). 2. T790M-like mutations in the hydrophobic core (T790M-like). 3. Insertions in the loop at the C-terminal end of the αC-helix in exon 20 (Ex20ins-L). 4. Mutations on the interior surface of the ATP-binding pocket or C-terminal end of the αC-helix, which is predicted to be P-loop and αC-helix compressing (PACC). We leverage two of the most influential subgroups (T790M-like and Ex20ins-L) to validate the performance of UdanDTI.

Besides, we identified 1,125 experimental drug-protein complex structures from the PDB (Protein Data Bank, https://www.rcsb.org/), which are included in the BindingDB dataset. They were selected as the test set to quantitatively compare interpretability. Their PDB IDs are listed in Table S4.

**Table S3.**
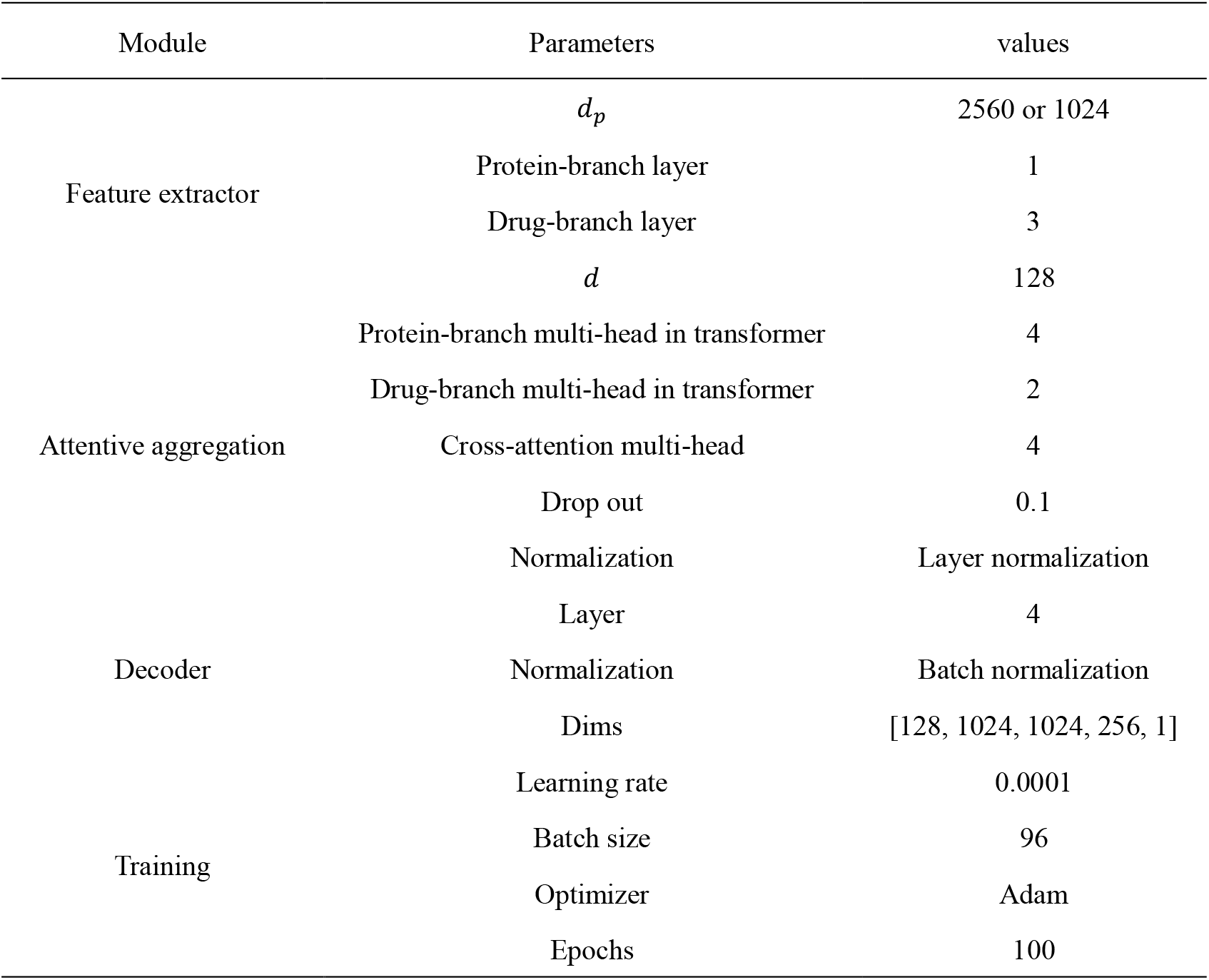
Hyperparameter setting.

**Table S4.**
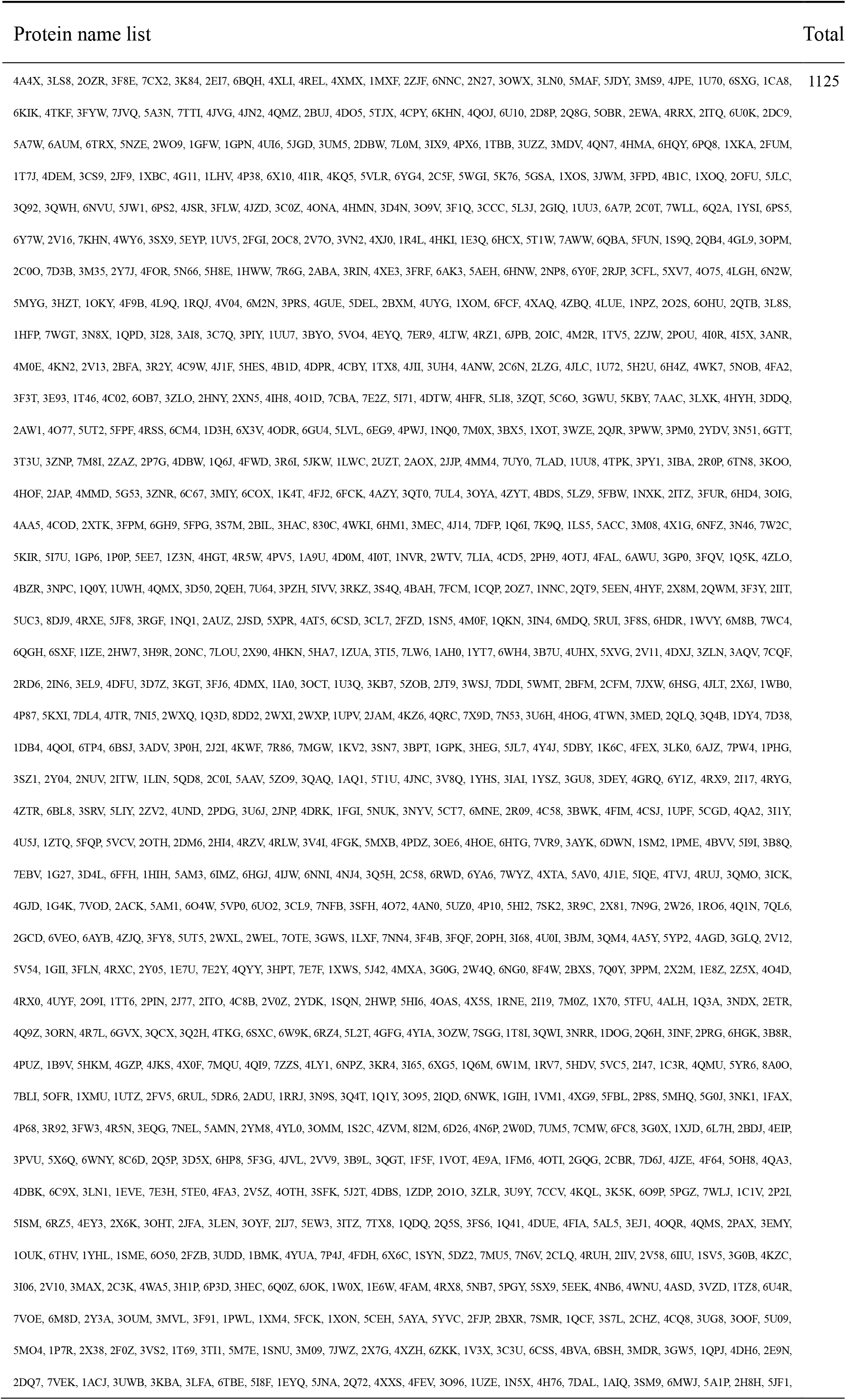
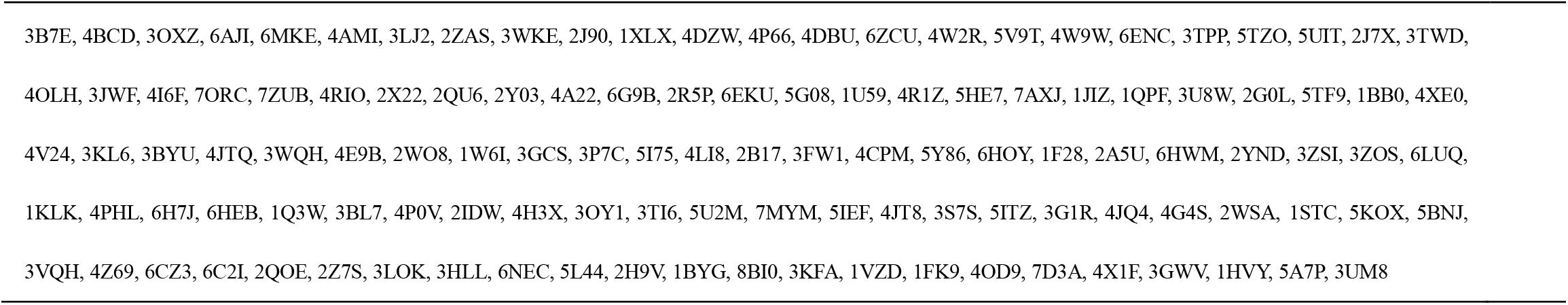
PDB ids of complexes selected.

## 2 Baselines

To evaluate UdanDTI comprehensively, we compared its performance against those of several state-of-the-art deep learning models renowned for their success in drug-target interaction (DTI) prediction.

1. RF employs the random forest algorithm on concatenated drug and protein feature vectors. We utilize ECFP4 fingerprinting[7] for drugs and PSC features[8] for proteins, a combination that proved to be highly effective.
2. DeepConv-DTI[9] utilizes CNNs and a global max-pooling layer to extract local patterns from protein sequences while applying a fully connected layer to the ECFP4 fingerprint of drugs. We follow the hyperparameter settings as described in the original paper.
3. GraphDTA[10] utilizes GNNs to encode drug molecular graphs and CNNs to encode protein sequences. A sigmoid function layer is added to the last fully connected layer with the cross-entropy loss function to adapt this regression model to a binary classification task.
4. MolTrans[4] leverages the transformer architecture to encode protein and drug sequences into feature embeddings, forming an interaction matrix. It subsequently applies CNNs and Fully Connected Networks (FCNs) to this interaction matrix for predictions. MolTrans output is also used to calculate the Top-N hit when comparing interpretability.
5. MGraphDTA[11] is motivated by the premise that shallow-layer GNNs struggle to learn complex molecule sub-structures like rings. It employs a hyper-deep GNN and a hyper-deep CNN to capture global representations of drugs and proteins, respectively. Shortcut connections in each layer mitigate over-smoothing.
6. MCANet[12] applies a multi-head cross-attention mechanism that shares weights between drug and protein representations. The key values of protein and drug feature maps are exchanged during multi-head attention, enhancing their feature representation, and both attention modules share the same weights. The MCANet model is used to calculate the Top-N hit for interpretability comparison.
7. DrugBAN[13] incorporates a bilinear attention network to fuse drug and protein representations decoded via a fully connected layer. It explicitly addresses interpretability by visualizing the pairwise bilinear attention map. Moreover, it employs a conditional domain adversarial network (CDAN) for knowledge transfer between the source and target domains to enhance cross-domain generalization. Vanilla DrugBAN is used to calculate the Top-N hit for interpretability comparison, and DrugBAN with a CDAN module is validated under the crossdomain setting.

These baselines were selected to represent a wide spectrum of deep learning methodologies in the DTI prediction domain, ranging from traditional architectures to recent advancements, encompassing interpretability and cross-domain generalization.

## 3 Sensitivity analysis

### 3.1 Under different hyperparameters

Figure S1 illustrates the AUROC performance of UdanDTI on the BioSNAP dataset under various hyperparameter settings. We investigated the impact of the learning rate, batch size, embedding dimension, and the number of attentive aggregation layers on UdanDTI. The experimental results indicate that UdanDTI’s performance does not exhibit significant differences after convergence (approximately 40 epochs), except for the potential underfitting due to the large learning rate.

**Figure S1.**
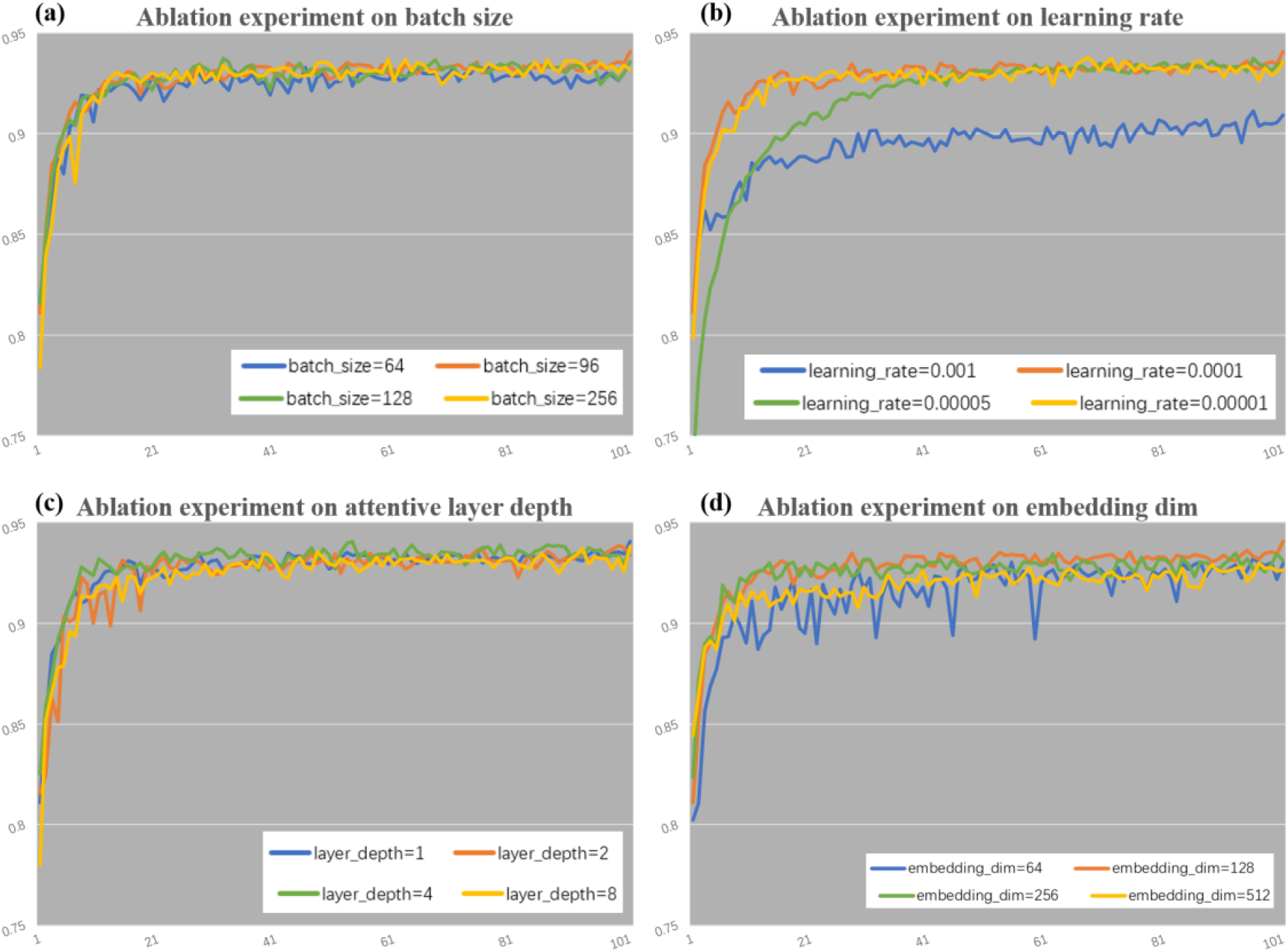
Learning curves (AUROC) with the different choices of hyperparameters on the BioSNAP dataset.

### 3.2 Across different protein family

We investigated UdanDTI’s performance across different protein families through experiments. Based on prior work, we utilized the GtoPdb database to map the test sets of BindingDB and BioSNAP into six major protein families: catalytic receptors, enzymes, G-protein-coupled receptors (GPCRs), ion channels, nuclear hormone receptors (NHRs) and transporters. Table S5 provides the statistics of six test subsets. The performance of UdanDTI across different protein families is illustrated in Table S6. Despite inherent biases within the datasets (such as inferior information for the transporters and NHRs family), UdanDTI maintains an overall high precision. In some critical protein families, notably GPCRs and ion channels, UdanDTI demonstrates more accurate predictions.

**Table S5.** Statistics across different protein family in test set.

| Dataset | Protein family | Drugs | Proteins | Interactions |
| --- | --- | --- | --- | --- |
| BioSNAP | GPCRs | 393 | 113 | 533 |
|  | Ion channels | 329 | 128 | 528 |
|  | NHRs | 132 | 29 | 139 |
|  | Catalytic receptors | 143 | 67 | 150 |
|  | Enzymes | 1744 | 699 | 2412 |
|  | Transporters | 215 | 100 | 240 |
| BindingDB | GPCRs | 574 | 188 | 707 |
|  | Ion channels | 466 | 65 | 522 |
|  | NHRs | 149 | 29 | 168 |
|  | Catalytic receptors | 580 | 63 | 747 |
|  | Enzymes | 4882 | 801 | 6618 |
|  | Transporters | 402 | 63 | 514 |

**Table S6.** Performance of UdanDTI across different protein family.

| Dataset | Protein family | AUROC | AUPRC |
| --- | --- | --- | --- |
| BioSNAP | GPCRs | 0.976±0.004 | 0.990±0.002 |
|  | Ion channels | 0.970±0.002 | 0.985±0.001 |
|  | NHRs | 0.933±0.004 | 0.962±0.003 |
|  | Catalytic receptors | 0.872±0.009 | 0.834±0.007 |
|  | Enzymes | 0.933±0.001 | 0.950±0.002 |
|  | Transporters | 0.912±0.008 | 0.894±0.008 |
| BindingDB | GPCRs | 0.945±0.003 | 0.969±0.002 |
|  | Ion channels | 0.973±0.003 | 0.940±0.001 |
|  | NHRs | 0.889±0.011 | 0.896±0.007 |
|  | Catalytic receptors | 0.946±0.005 | 0.945±0.004 |
|  | Enzymes | 0.966±0.001 | 0.953±0.001 |
|  | Transporters | 0.981±0.004 | 0.963±0.006 |

### 3.3 5-fold and 10-fold cross validation results

**Table S7.** 5-fold and 10-fold cross validation of UdanDTI.

| Dataset | AUROC | AUPRC | AUROC<sub>mean</sub> | AUROC<sub>std</sub> | AUPRC<sub>mean</sub> | AUPRC<sub>std</sub> |
| --- | --- | --- | --- | --- | --- | --- |
| BioSNAP<sub>5-fold</sub> | 0.940±0.004 | 0.944±0.005 | 0.941 | 0.0028 | 0.944 | 0.0034 |
| BioSNAP<sub>10-fold</sub> | 0.942±0.004 | 0.943±0.006 | 0.942 | 0.0029 | 0.944 | 0.0038 |
| BindingDB<sub>5-fold</sub> | 0.967±0.001 | 0.954±0.002 | 0.967 | 0.0007 | 0.953 | 0.0014 |
| BindingDB<sub>10-fold</sub> | 0.967±0.002 | 0.955±0.004 | 0.967 | 0.001 | 0.954 | 0.0022 |

## 4 Debiasing evaluation

To further evaluate the degree of overfitting of the DTI models and emphasize the drug-bias trap, we modified the test set of BioSNAP. Specifically, we randomly selected 16 drugs and 200 target proteins in the test set and predicted them with UdanDTI, DrugBAN, MCANet, and MolTrans, respectively. We then follow the YUEL [14] model to replace each protein sequence with a single Alanine. Individual Alanine should not have a strong binding affinity with drugs, forcing DTI models to use only drug information to predict. Visualizing the test results revealed that all other models except UdanDTI failed to adequately distinguish the variability among target proteins. In other words, existing DTI models tend to make drug-centric predictions.

**Table S8.**
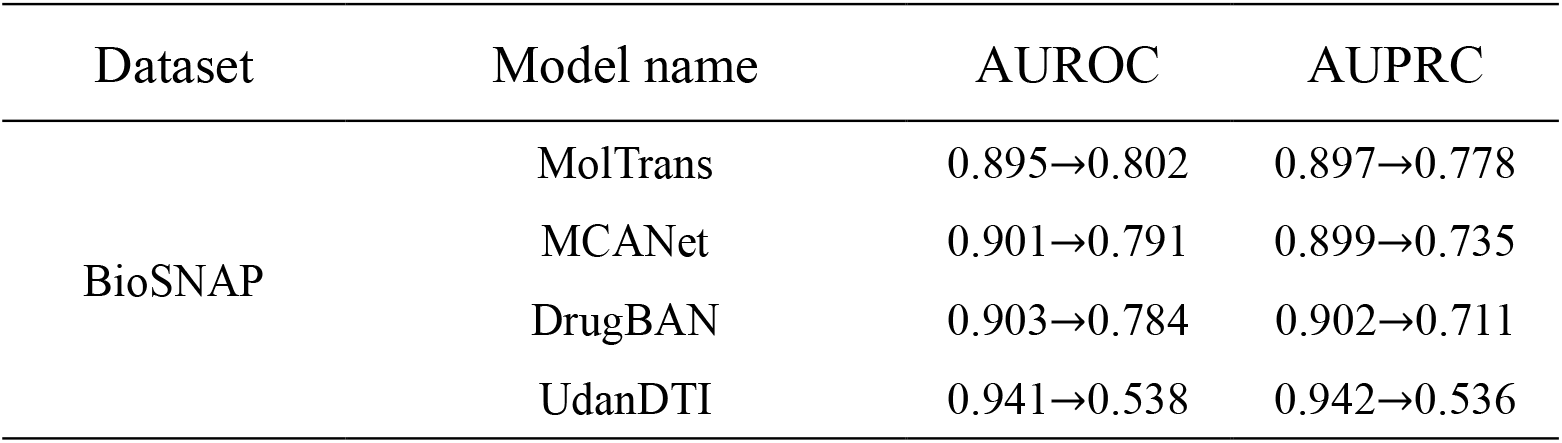
Debiasing evaluation on BioSNAP.

| Dataset | Model name | AUROC | AUPRC |
| --- | --- | --- | --- |
| BioSNAP | MolTrans | 0.895→0.802 | 0.897→0.778 |
|  | MCANet | 0.901→0.791 | 0.899→0.735 |
|  | DrugBAN | 0.903→0.784 | 0.902→0.711 |
|  | UdanDTI | 0.941→0.538 | 0.942→0.536 |

## 5 Detailed comparison of interpretability on protein

To further evaluate the interpretability on protein sequence, we further compared the frequency of effective hits of different amino acids by each DTI model.

**Table S9.** Frequency of effective hits.

| Amino acid type | MolTrans (%) | MCANet (%) | DrugBAN (%) | UdanDTI (%) |
| --- | --- | --- | --- | --- |
| GLY | 6.27 | 6.22 | 6.24 | 5.03 |
| CYS | 1.73 | 2.13 | 2.21 | 1.62 |
| LYS | 7.17 | 6.44 | 7.27 | 5.67 |
| PHE | 5.73 | 6.51 | 5.64 | 7.61 |
| ILE | 5.36 | 7.98 | 6.83 | 6.37 |
| LEU | 9.65 | 9.46 | 9.6 | 11.42 |
| ASN | 3.71 | 3.32 | 4.54 | 4.92 |
| ARG | 6.7 | 5.59 | 5.72 | 5.41 |
| GLU | 6.31 | 5.09 | 5.02 | 5.6 |
| MET | 2.46 | 2.39 | 2.6 | 3.17 |
| ALA | 5.71 | 4.99 | 5.36 | 3.97 |
| TYR | 5.05 | 4.95 | 6.19 | 6.83 |
| HIS | 2.53 | 2.42 | 3.34 | 4.39 |
| TRP | 2.64 | 3.13 | 2.79 | 3.03 |
| GLN | 3.7 | 3.89 | 4.02 | 3.16 |
| PRO | 2.85 | 3.3 | 3.11 | 2.49 |
| VAL | 8.32 | 9.25 | 7.54 | 5.76 |
| THR | 4.27 | 4.34 | 3.88 | 3.67 |
| SER | 5.02 | 4.03 | 4.01 | 4.55 |
| ASP | 4.83 | 4.57 | 4.09 | 5.34 |

## 6 Comparison under some other splitting

For the cross-domain splitting strategy, we followed the previous work [13]. Specifically, we use the binarized ECFP4 feature to represent drug compounds and the integral PSC feature to represent target proteins. Then, we use the single-linkage clustering method to cluster them separately. We randomly select 60% drug clusters and 60% protein clusters from the clustering results and regard all associated drug-target pairs with them as source domain data. The associated pairs in the remaining clusters are considered to be source domain data. In fact, clustering protein and drug feature representations, respectively, can simultaneously mitigate potential biases in both drugs and proteins.

To comprehensively assess the scalability of models, we partitioned the large-scale datasets, BioSNAP and BindingDB, using three other splitting methods: unseen drugs, unseen proteins, and sphere exclusion clustering. Respectively, we randomly selected 10% of the drugs or proteins and their associated drug-protein pairs to form the test set. The remaining data was used to train the models. Besides, we employed ECFP4 to generate molecular features for drug molecules and assigned all compounds into different clusters using the algorithm recommended in the paper [15]. Finally, we randomly selected drug-protein pairs corresponding to 80% of the drug clusters as the training set.

We replicated and re-trained four advanced models (RF, MolTrans[4], MCANet[12], and DrugBAN[13]). The comparative results are shown in Table S9, UdanDTI still exhibits the best performance.

**Table S10.**
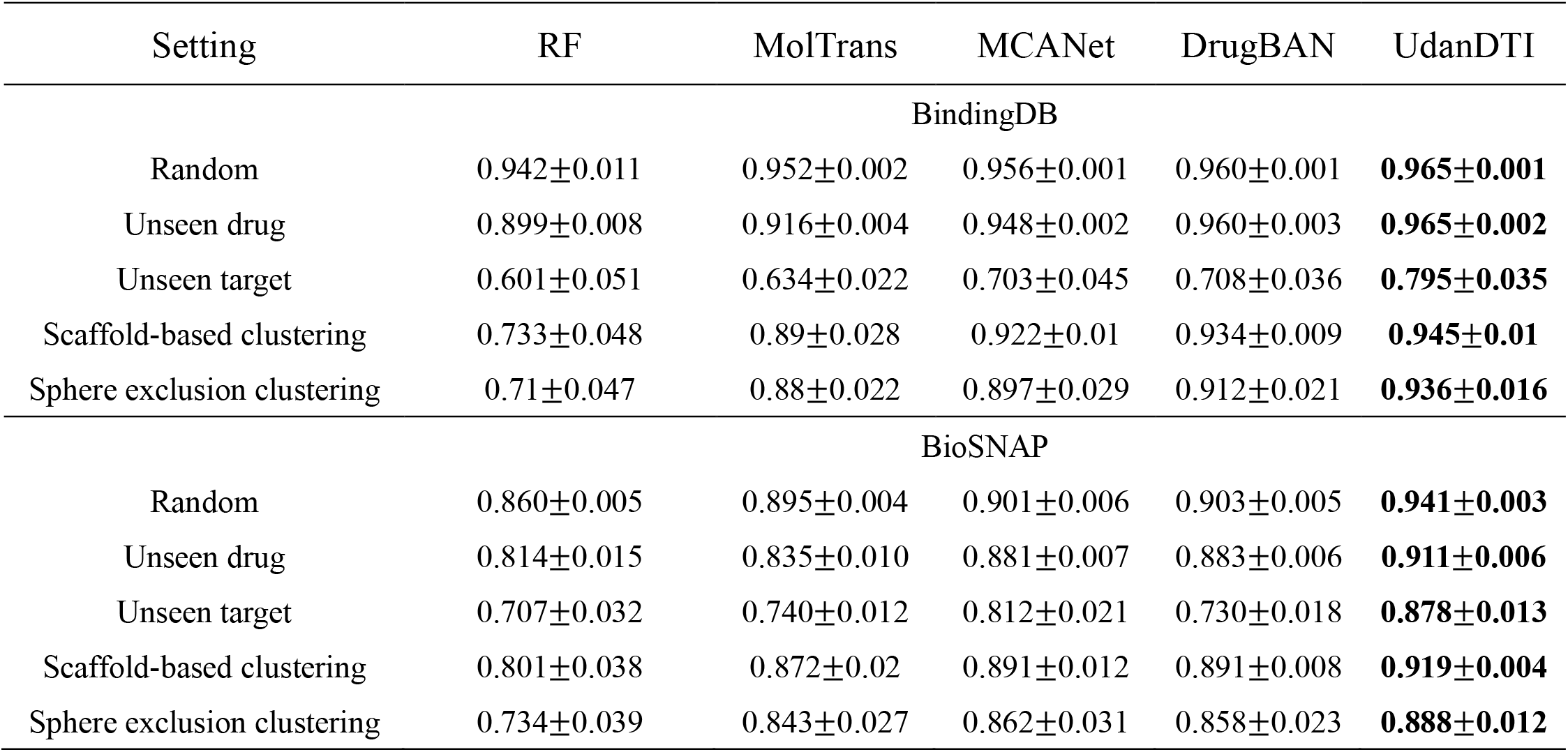
Comparison (AUROC) under the setting of unseen drugs/proteins.

| Setting | RF | MolTrans | MCANet | DrugBAN | UdanDTI |
| --- | --- | --- | --- | --- | --- |
| BindingDB |  |  |  |  |  |
| Random | 0.942±0.011 | 0.952±0.002 | 0.956±0.001 | 0.960±0.001 | **0.965±0.001** |
| Unseen drug | 0.899±0.008 | 0.916±0.004 | 0.948±0.002 | 0.960±0.003 | **0.965±0.002** |
| Unseen target | 0.601±0.051 | 0.634±0.022 | 0.703±0.045 | 0.708±0.036 | **0.795±0.035** |
| Scaffold-based clustering | 0.733±0.048 | 0.89±0.028 | 0.922±0.01 | 0.934±0.009 | **0.945±0.01** |
| Sphere exclusion clustering | 0.71±0.047 | 0.88±0.022 | 0.897±0.029 | 0.912±0.021 | **0.936±0.016** |
| BioSNAP |  |  |  |  |  |
| Random | 0.860±0.005 | 0.895±0.004 | 0.901±0.006 | 0.903±0.005 | **0.941±0.003** |
| Unseen drug | 0.814±0.015 | 0.835±0.010 | 0.881±0.007 | 0.883±0.006 | **0.911±0.006** |
| Unseen target | 0.707±0.032 | 0.740±0.012 | 0.812±0.021 | 0.730±0.018 | **0.878±0.013** |
| Scaffold-based clustering | 0.801±0.038 | 0.872±0.02 | 0.891±0.012 | 0.891±0.008 | **0.919±0.004** |
| Sphere exclusion clustering | 0.734±0.039 | 0.843±0.027 | 0.862±0.031 | 0.858±0.023 | **0.888±0.012** |

## 7 Ablation experiments

### 7.1 Ablation experiments about LLMs

We completed the ablation experiments on LLMs in the table below. We compared the results of four protein LLMs (ProGen[16], Prose[17], ESM[18], and ProtBert[19]) and two drug molecule LLMs (ChemBerta[20] and Molformer[21]), as well as their possible combinations, on the BioSNAP dataset. Obviously, the non-balanced dual-branch model using only ESM or ProtBert protein LLM is the best.

ProtBert^1^ and ProtBert^2^ are LLM with different weight versions (ProtT5XLUniref50 and ProtBert-BFD respectively).

**Table S11.** Ablation results about LLMs on BioSNAP.

| Dual-branch network | LLM | AUROC | AUPRC |
| --- | --- | --- | --- |
| Balanced | – | 0.887±0.004 | 0.901±0.005 |
| Unbalanced<br>(drug-oriented) | ChemBertA | 0.894±0.004 | 0.908±0.004 |
|  | Molformer | 0.885±0.007 | 0.893±0.011 |
| Unbalanced<br>(protein-oriented) | ProtBert<sup>1</sup> | **0.941±0.003** | 0.942±0.004 |
|  | ProtBert<sup>2</sup> | 0.926±0.002 | 0.923±0.002 |
|  | ESM | 0.937±0.003 | **0.943±0.002** |
|  | ProGen | 0.924±0.005 | 0.928±0.007 |
|  | Prose | 0.935±0.003 | 0.937±0.005 |
| Balanced | ESM + ChemBertA | 0.934±0.004 | 0.932±0.006 |
|  | ESM + Molformer | 0.901±0.007 | 0.900±0.008 |
|  | ProtBert<sup>1</sup> + ChemBertA | 0.932±0.004 | 0.934±0.007 |
|  | ProtBert<sup>1</sup> + Molformer | 0.907±0.005 | 0.905±0.005 |

ProtBert<sup>1</sup> and ProtBert<sup>2</sup> are LLM with different weight versions (ProtT5XLUniref50 and ProtBert-BFD respectively).

### 7.2 Ablation experiments about attentive aggregation module

We completed the ablation experiments on the attentive aggregation module in the table below. For a fair comparison, we employed various feature fusion modules into the same non-balanced dual-branch model on the BioSNAP dataset. The concatenation means combining the representations of drug and protein, and the cross-attention means directly incorporating all protein and drug information into the cross-attention mechanism. The bilinear attention means using the bilinear attention network from previous work [22]. Moreover, we construct a new baseline using a simple layer to learn the dynamic weights for summing the head representations of protein and drug representations in UdanDTI.

The experiments demonstrated the superiority of our attentive aggregation module.

**Table S12.** Ablation results about feature fusion modules on BioSNAP.

| Fusion module | AUROC | AUPRC |
| --- | --- | --- |
| Concatenate | 0.904±0.007 | 0.902±0.008 |
| Cross attention | 0.930±0.005 | 0.933±0.004 |
| Bilinear attention | 0.927±0.005 | 0.925±0.003 |
| Dynamic weights | **0.943±0.006** | 0.940±0.007 |
| Attentive aggregation | 0.941±0.003 | **0.942±0.004** |

### 7.3 Ablation experiments about representations of protein

We completed the ablation experiments on the representation of protein in the table below. We constructed two extra baselines using indexical one-hot encoding coding (to indicate the type of amino acid) and BLOSUM62 matrix encoding, respectively, demonstrating the advantages of LLM with deep layers in capturing the structural properties of proteins.

**Table S13.**
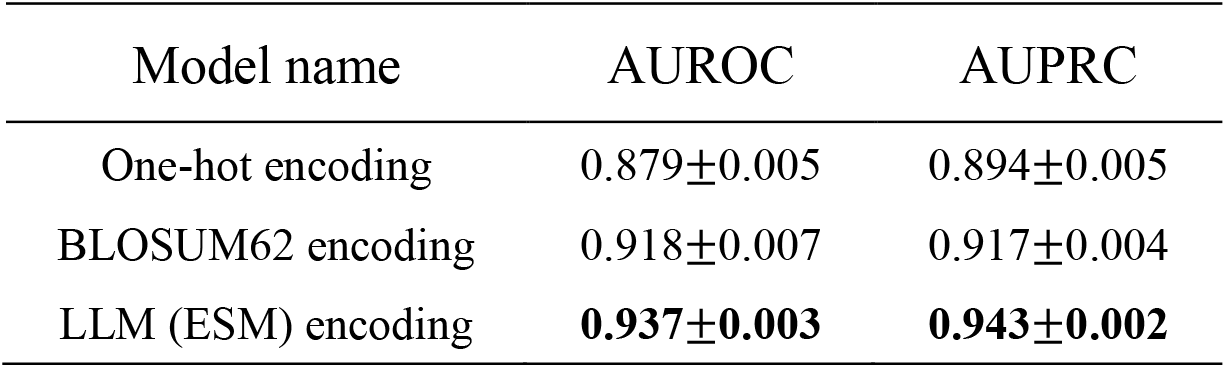
Ablation results about representations of protein on BioSNAP.

| Model name | AUROC | AUPRC |
| --- | --- | --- |
| One-hot encoding | 0.879±0.005 | 0.894±0.005 |
| BLOSUM62 encoding | 0.918±0.007 | 0.917±0.004 |
| LLM (ESM) encoding | **0.937±0.003** | **0.943±0.002** |

## 8 Heatmap between amino acids and functional groups

To further elucidate the working mechanism of UdanDTI, we conducted an analysis of predictions for 1,125 structures within the BindingDB dataset and compared them against the experimental results (structures). For each DTI pair, we recorded the attentive amino acids identified by UdanDTI, along with the amino acids around the protein pockets (within 6 Å) in the actual structures. Additionally, we employed RDKit to record the functional groups involved in the reaction. Figure S2 illustrates the heatmaps of model predictions and experimental statistics. It demonstrates that UdanDTI effectively mitigates the drug-bias trap, indicating a genuine understanding of potential interaction patterns.

**Figure S2.**
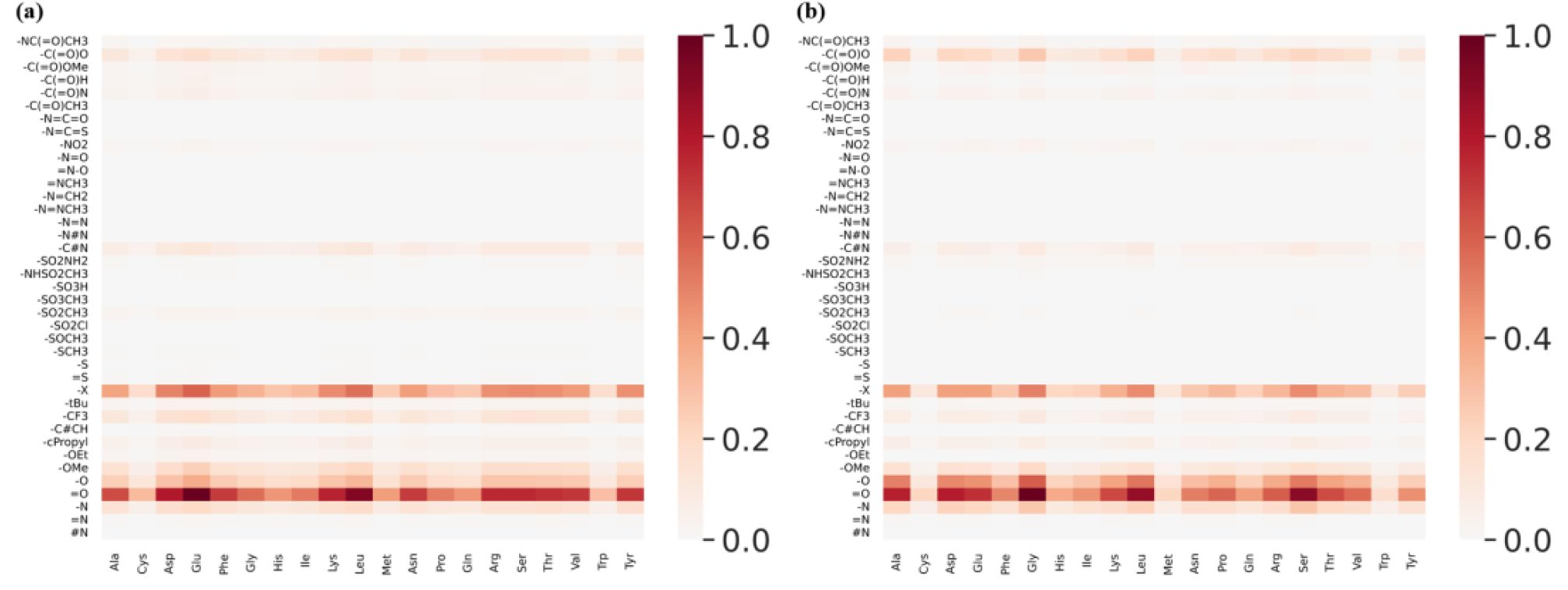
Comparison heatmaps of model predictions (a) and experimental statistics (b).

## 9 Comparison with DiffDock

Here, we emphasize the superior performance of UdanDTI over advanced docking models like DiffDock[23] in virtual screening. Virtual screening aims to identify molecules within a molecular database that might bind to a target protein rather than predicting precise binding poses, making it more relevant to real-world scenarios. As depicted in Figure S3, when performing high-throughput screening on the same target protein across different drug molecular databases, the speed advantage of UdanDTI becomes increasingly prominent as the data size grows. When dealing with database numbering in the billions, the running time of DiffDock becomes intolerable. Furthermore, if pre-generating corresponding protein embeddings using LLM and storing them on disk, UdanDTI’s speed can be further enhanced.

**Figure S3.**
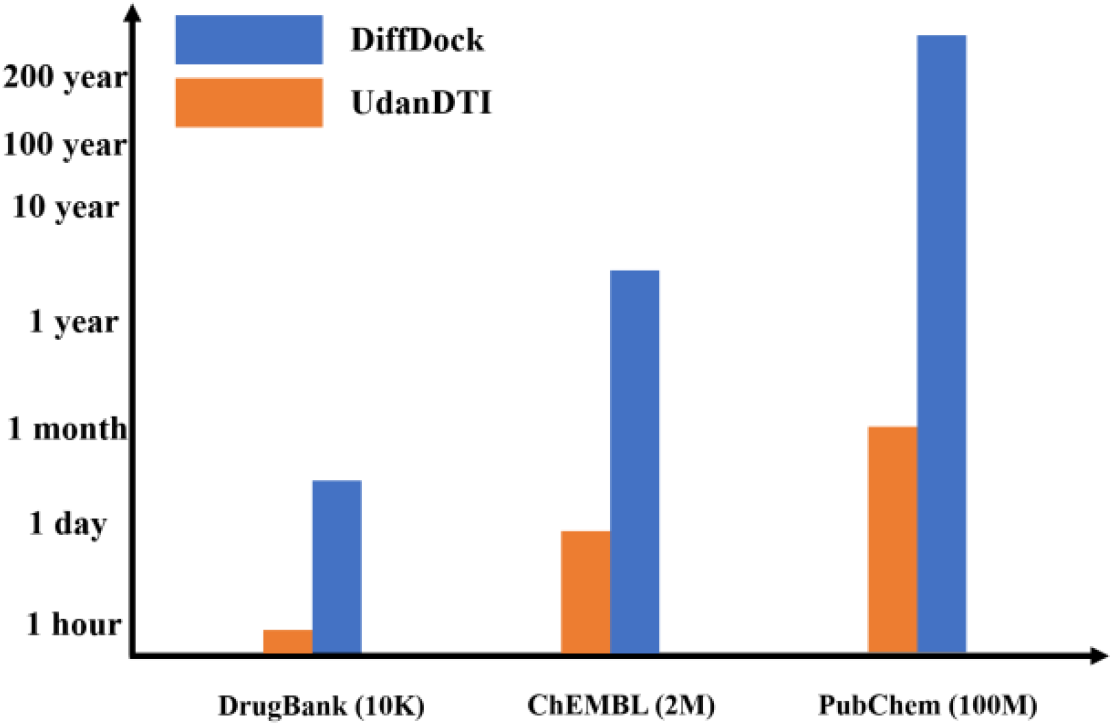
Time analysis between UdanDTI and DiffDock in virtual screening.

Moreover, UdanDTI’s accuracy in virtual screening is significantly higher than that of DiffDock. We selected five target proteins (named Q6V1X1, P10635, P48635, Q86TI2, and P63316 in UniProt dababase[24]) and removed relevant drug-target pairs from UdanDTI’s training set, simulating screening among 14,643 drugs from the BindingDB. For each target protein, we compared the top 50 drugs with the highest confidence scores from both UdanDTI and DiffDock, respectively. As shown in Table S13, UdanDTI notably provides researchers with a greater number of genuinely interactive drug candidates.

**Table S14.** Number of interactive drugs in top 50 candidates provided by UdanDTI and DiffDock.

| Model name | Q6V1X1 | P10635 | P48635 | Q86TI2 | P63316 |
| --- | --- | --- | --- | --- | --- |
| DiffDock | 3 | 4 | 0 | 2 | 5 |
| UdanDTI | 19 | 19 | 18 | 33 | 34 |

## 10 Implementation and Experimental Environment

### Model Parameters

UdanDTI was implemented in Python 3.8. The main network architecture was implemented in PyTorch 2.0, while the data extraction and processing were written by RDKit 2023.3.1, DGL 1.1.1, and DGL-lifeSci 0.3.2 separately. ESM and ProtBert in huggingface (https://huggingface.co/) were used for amino acid sequence encoding, and all pretrained parameters were frozen. We chose the esm2_t36_3B_UR50D and prot_bert_bfd versions. They used 36 and 30 transformer layers separately, with the 2560-D and 1024-D protein embeddings. The GCN-based drug encoder is set to 3 layers with the same 128-D hidden dimensions. The maximum length of the sequence input to the encoder was 290 for drugs and 1200 for proteins, accounting for 98% of the entire dataset. In the mutual information learning module, two parallel branches had the same architecture, concatenating 1 transformer layer and 1 cross-attention layer. However, we set a 4-head transformer in the protein branch and a 2-head transformer in the drug branch. The attention heads in crossattention were both set to 4. Besides, the dropout rate of the interaction layer was set to 0.1, and the batch size was set to 128. The optimizer used Adam, and the learning rate was set to 0.0001.

### Hardware

1 NVIDIA A40 Tensor Core GPU was used for training and testing.

## Notes

### Competing Interest Statement

The authors have declared no competing interest.

